# Differential effects of aging, Alzheimer’s pathology, and *APOE4* on longitudinal functional connectivity and episodic memory in older adults

**DOI:** 10.1101/2024.12.11.627967

**Authors:** Larissa Fischer, Jenna N. Adams, Eóin N. Molloy, Niklas Vockert, Jennifer Tremblay-Mercier, Jordana Remz, Alexa Pichet Binette, Sylvia Villeneuve, PREVENT-AD Research Group, Anne Maass

## Abstract

**INTRODUCTION:** Both aging and Alzheimer’s disease (AD) affect episodic memory networks. How this relates to region-specific early differences in functional connectivity (FC), however, remains unclear.

**METHODS:** We assessed resting-state FC strength in the medial temporal lobe (MTL) - posteromedial cortex (PMC) - prefrontal network and cognition over two years in cognitively normal older adults from the PREVENT-AD cohort.

**RESULTS:** FC strength within PMC and between posterior hippocampus and inferomedial precuneus decreased in “normal” aging (amyloid- and tau-negative adults). Lower FC strength within PMC was associated with poorer longitudinal episodic memory performance. Increasing FC between anterior hippocampus and superior precuneus was related to higher baseline AD pathology. Higher FC strength was differentially associated with memory trajectories depending on *APOE4* genotype.

**DISCUSSION:** Findings suggest differential effects of aging and AD pathology on longitudinal FC. MTL-PMC hypoconnectivity was related to aging and cognitive decline. Furthermore, MTL-PMC hyperconnectivity was related to early AD pathology and cognitive decline in *APOE4* carriers.

**Graphical abstract.**
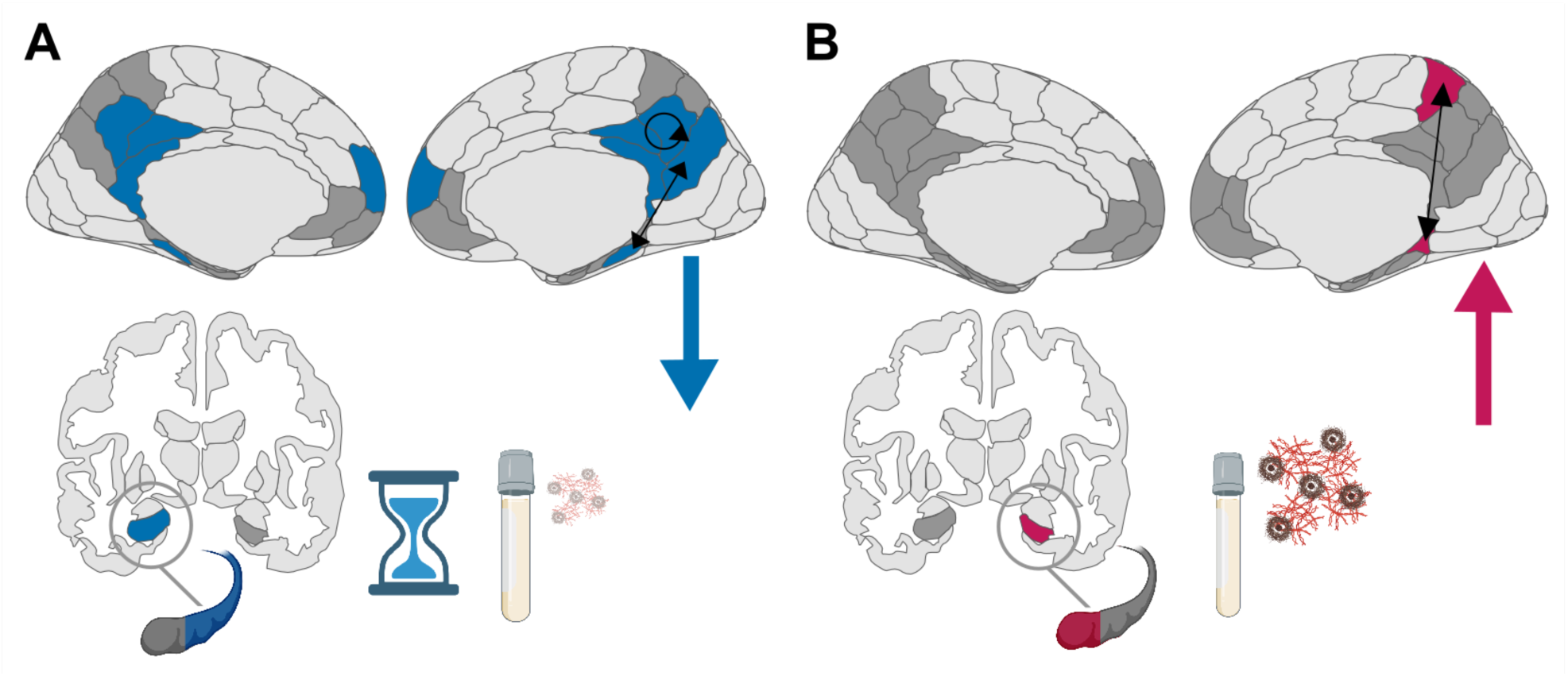
**A)** “Normal aging” is characterized by a longitudinal decrease in functional connectivity. **B)** Cognitively unimpaired older adults with more Alzheimer’s pathology at baseline (measured via cerebrospinal fluid) exhibit a longitudinal increase in functional connectivity.

## 1. Background

Episodic memory deficits in aging and Alzheimer’s disease (AD) are hypothesised to partly result from disruptions in medial temporal lobe (MTL) and neocortical episodic-memory dependent networks.^2,3^ Although intact episodic memory is key for independent living, it is among the first cognitive functions to deteriorate with age and AD.^4^ Functional connectivity (FC) of the MTL, the posteromedial cortex (PMC) and medial prefrontal cortex is crucial for remembering prior experiences.^5,6^ Changes in FC with aging and AD are, however, complex and poorly understood. Differentiating the respective functional dynamics offers significant potential for early identification of individuals in (future) need of clinical intervention and for targeting aberrant FC processes as intervention pathways.^7,8^

AD pathology consists of amyloid-beta (Aβ) plaques and tau tangles, starting in the PMC^9^ and MTL^10^, respectively, and is present in a substantial proportion of cognitively normal older adults.^11,12^ Aberrant functional network measures have been associated, predominantly cross-sectionally, with Aβ^13–15^ and tau^16–18^ in this population. More specifically, animal models^19,20^ and human studies^15,17,21–23^ suggest that amyloid is rather related to hyperconnectivity, while tau may be first related to hyper- and then hypoconnectivity in later AD stages. While animal models can provide direct evidence from cell recordings, aberrant FC in functional magnetic resonance imaging (fMRI) studies is observed indirectly via lower co-variation in blood oxygen level-dependent (BOLD) signal between regions.

Pinning down brain areas that show the earliest FC changes with AD pathology remains a challenge. Regarding the MTL, cross-sectional studies suggested that the parahippocampal gyrus and hippocampus are among the first MTL regions that show functional alterations with early AD pathology.^24,25^ Given their role as hub regions within the MTL-PMC network, these changes align with the hypothesis of hub hyperconnectivity in response to adverse influences.^26^ Further, especially the anterior hippocampus, entorhinal and perirhinal cortex seem to be involved in early aberrant amyloid-related hyperconnectivity.^27–29^ Regarding “normal” aging, hypoconnectivity has been cross-sectionally described in amyloid-negative individuals.^17,28^ Most longitudinal studies on “normal” aging did, however, not assess AD pathology or AD-risk factors like the *APOE4* genotype, leaving significant uncertainty about the effects of aging in the absence of Aβ and tau.

As a result, it remains unclear which specific age-related and AD-related changes in FC can be distinguished, and how these changes relate to episodic memory performance. Most studies to date were limited to cross-sectional data and few findings on longitudinal functional and cognitive changes including information for both Aβ and tau measures combined have been reported.^30^ Longitudinal imaging studies are crucial to advance the understanding of aging and AD-related processes over time, especially when evaluating cognitive trajectories.^31,32^

To disentangle FC changes related to “normal” aging and early neurodegenerative pathogenesis, we quantified longitudinal changes in resting-state FC (rsFC) in the MTL-PMC- prefrontal network in a group of cognitively normal older adults (a) without evidence of AD pathology and (b) with available longitudinal AD biomarker data. Specifically, we aimed to relate rsFC changes to aging, Aβ and tau pathology, *APOE4* genotype, and longitudinal memory performance in the PREVENT-AD cohort^33^ of cognitively unimpaired older adults with this preregistered study.^1^

We hypothesized for the investigated memory network that i) decreasing rsFC strength over time, especially with older age, is visible in the presence of low pathology (A^-^T^-^), ii) increasing rsFC is associated with higher early AD pathology, especially in *APOE4* carriers, iii) decreasing integration of hub regions and meaningful subnetwork segregation is visible over time, especially with older age, with AD pathology and in *APOE4* carriers, iv) mild age-related (A^-^T^-^) decline or less practice effects in episodic memory performance are associated with steeper decrease in rsFC strength over time, especially with older age, v) higher or increasing rsFC strength could be an initial beneficial or compensatory process if predicting maintained episodic memory performance or a detrimental process if predicting decline in performance.

## 2. Methods

### 2.1. Sample and study design

All participants were cognitively normal older adults from the Pre-symptomatic Evaluation of Experimental or Novel Treatments for Alzheimer’s Disease (PREVENT-AD) cohort.^33,34^ PREVENT-AD is an ongoing longitudinal study with first enrollments starting in 2011. The data we used were part of data release 7.0 with parts of the data being publicly available. Participants self-reported having a parental history of AD dementia or two siblings diagnosed with AD dementia and were above 60 years of age at enrollment. Participants aged between 55 and 59 were also included if their age was less than 15 years away from their affected relative’s age at onset. Participants had no major neurological or psychiatric illnesses at baseline and were screened for normal cognition with the Montreal Cognitive Assessment (MoCA) questionnaire and the Clinical Dementia Rating (CDR) Scale. If a participant’s cognitive status was in doubt (MoCA < 27 or CDR > 0), the participant had to undergo exhaustive neuropsychological evaluation and was excluded upon a diagnosis of probable MCI.

We formed two subsamples to answer our specific research questions of dissociable effects of aging and AD pathology. Subsample A (“normal” aging sample) consisted of Aβ- and tau-negative older adults (A^-^T^-^, N=100, 63±6years, 72 female, 30 *APOE4* carriers) according to their cerebrospinal fluid (CSF) or positron emission tomography (PET) data. Subsample B (AD-biomarker sample) contained all available data from older adults with longitudinal CSF measurements (N=70, 63±5years, 48 female, 24 *APOE4*), independent of biomarker status. The participants included in subsample A completed an MRI scan at baseline and follow-up assessments after 12 (FU12) and 24 months (FU24). To define this sample of A^-^T^-^ older adults, we used PET or CSF data to evaluate biomarker status for Aβ (A^+/-^) and tau (T^+/-^). This allowed us to include more participants, as some participants had CSF but no PET measurements and vice versa. Biomarker positivity was evaluated based on the following research cut-offs defined by the PREVENT-AD Research Group: CSF Aβ_1-42_ < 850 pg/ml, CSF p-tau_181_ > 60 pg/ml, for entorhinal tau PET SUVR > 1.3, and for whole brain Aβ PET SUVR > 1.39. Participants had to be negative via CSF at the last time-point of MRI and cognitive assessment we investigated, which was FU24. If only PET was available, they had to be negative at this measurement, which took place on average 6 years after baseline. If PET and CSF for time-points later than 24 months were available, both had to be below the respective threshold. The participants included in subsample B had to have MRI and CSF measurements at baseline and at FU24 to enable longitudinal assessment. 39 participants were included (i.e. overlapping) in both subsamples.

Quality control after preprocessing led to exclusions of participants. Specifically, seven participants in subsample A and one participant in subsample B were excluded from further analysis due to > 20% of volumes flagged as outliers (see section 2.7 for details). This resulted in a final subsample A of 93 older adults (A^-^T^-^, 63±5years, 68 female, 28 *APOE4* carriers) and subsample B of 69 older adults (62±6years, 48 female, 23 *APOE4* carriers), see Table 1.

**Table 1:** Final sample demographics after exclusions.

| Subsample | N | Age in years | Education in years | Female sex | *APOE4* carrier | CSF Aβ<sub>1-42</sub> at baseline | CSF p-tau<sub>181</sub> at baseline | Aβ PET SUVR | Tau PET SUVR |
| --- | --- | --- | --- | --- | --- | --- | --- | --- | --- |
| A | 93 | 63.26<br>(4.69) | 15.91<br>(3.58) | 68<br>(73%) | 28<br>(30%) | N = 46<br>1191.96<br>(183.86) | N = 46<br>40.52<br>(14.64) | N = 78<br>1.174<br>(0.09) | N = 78<br>1.026<br>(0.09) |
| B | 69 | 62.71<br>(5.64) | 15.13<br>(2.97) | 48<br>(70%) | 23<br>(33%) | N = 69<br>1142.06<br>(264.85) | N = 69<br>47.25<br>(19.37) | N = 45<br>1.260<br>(0.22) | N = 45<br>1.032<br>(0.08) |

Mean (standard deviation) or number (percentage) of participants. 39 participants were included in both subsamples. All participants identified as either female or male. Two *APOE4* carriers in subsample A were homozygous; there were no homozygous *APOE4* carriers in subsample B. For tau PET, entorhinal tau was investigated. For Aβ PET, global Aβ was used. N = number. CSF = cerebrospinal fluid. Aβ = amyloid-beta. PET = positron emission tomography. SUVR = standardized uptake value ratio.

All study procedures and experimental protocols were approved by the McGill University Institutional Review Board and/or the Douglas Mental Health Institute Research Ethics Board. Written informed consent was obtained from all participants prior to each experimental procedure and they were financially compensated for their time. For an overview of the sessions, see Figure 1A.

**Fig. 1.**
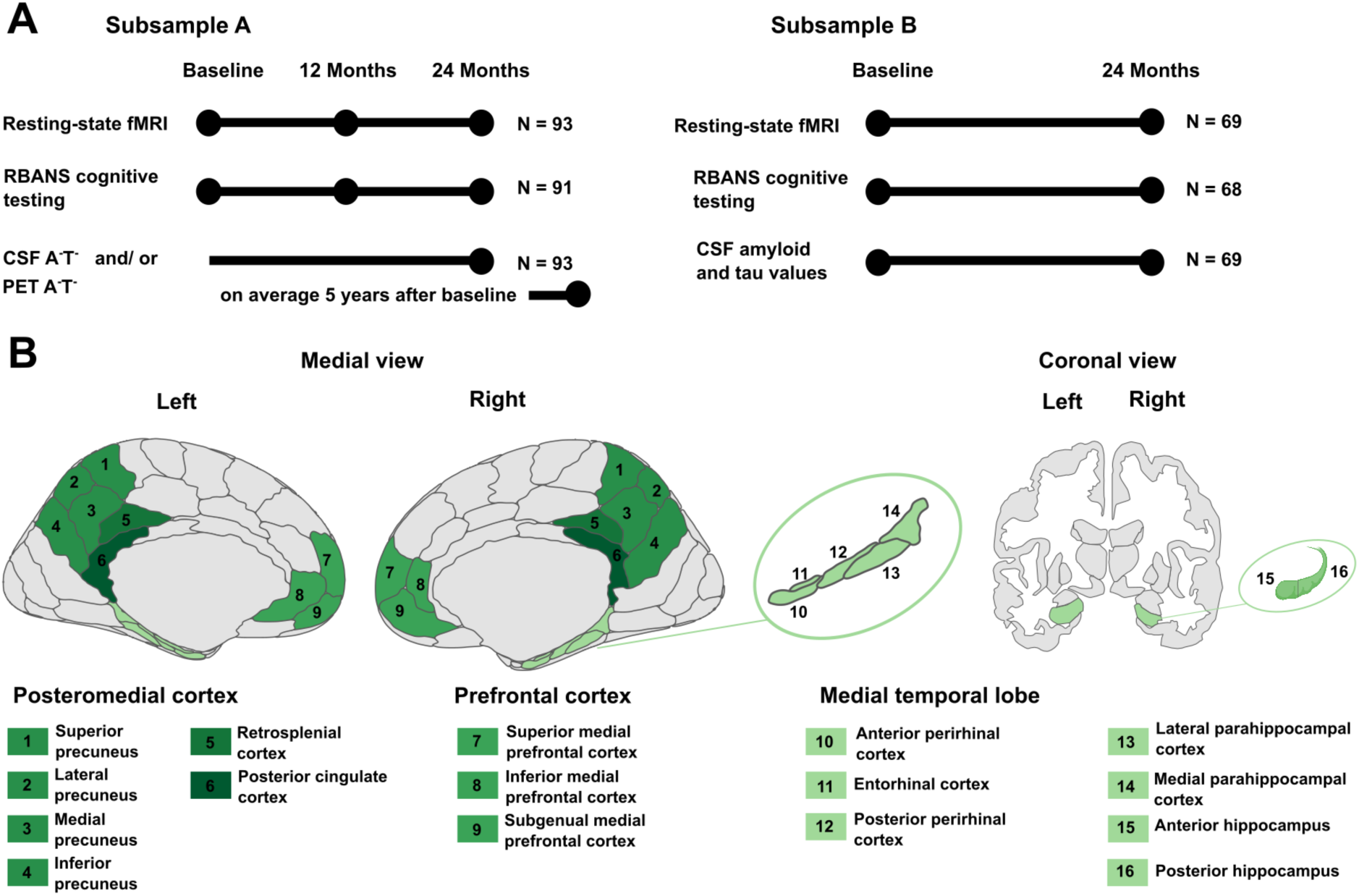
Study Design. Illustrations are presented in neurological view. Brain regions considered as a subregion of a larger region (precuneus, medial prefrontal cortex, and medial temporal lobe) are presented in the same shade of green. Retrosplenial cortex and posterior cingulate cortex were not further divided into subregions. **A)** Visualization of included sessions. Subsample A (“normal” aging) consisted of amyloid- and tau-negative (A^-^T^-^) older adults. Subsample B consisted of CSF-characterized older adults. **B)** Visualization of regions of interest. Medial view is based on the Brainnetome atlas, which was used for ROI definition. The coronal view is based on the aseg atlas for illustration of the hippocampus only. fMRI = functional magnetic resonance imaging. CSF = cerebrospinal fluid. PET = positron emission tomography. RBANS = Repeatable Battery for Assessment of Neuropsychological Status.

### 2.2. Cerebrospinal fluid collection

Participants who consented to this procedure contributed CSF samples from a lumbar puncture (LP) on a separate day around the main annual measurement, as described by Tremblay-Mercier and colleagues.^33^ The LP took place on average 28 days after the annual visit. A large-bore introducer was inserted at the L3-L4 or L4-L5 intervertebral space, then an atraumatic Sprotte 24 ga. spinal needle was used to puncture the dura. Up to 30 ml of CSF were withdrawn in 5.0 ml polypropylene syringes, centrifuged at room temperature for 10 min at ∼ 2000g, aliquoted in 0.5 ml polypropylene cryotubes, and quick-frozen at −80 °C for long-term storage. CSF measurements of interest for this study were Aβ_1-42_ and p-tau_181_ and the p-tau_181_/Aβ_1-42_ ratio to assess the effect of AD pathology. The ratio is an established measure in AD research^35^ and was included to assess the combined effect of amyloid-related tau hyperphosphorylation.

### 2.3. *APOE* genotyping

All included participants were genotyped for APOE via a QIASymphony apparatus. For details, see the description by Tremblay-Mercier and colleagues^33^. Participants who featured at least one copy of the *APOE4* allele were included in the carrier group while participants without an *APOE4* allele were allocated to the non-carrier group.

### 2.4. Assessment of episodic memory performance

Cognition was measured via the longitudinal Repeatable Battery for Assessment of Neuropsychological Status (RBANS) at each session.^36^ Different versions of the RBANS were used at each follow-up session to reduce practice effects. Our main measure of interest was the RBANS delayed memory index score, which integrates assessments from word-list recognition as well as delayed free recall of figures, stories, and word lists. To investigate the specificity of the associations between longitudinal memory performance and FC, we also used the attention index score as a memory-independent measure in a control analysis. The attention index score consists of two digit span subtests. RBANS data from baseline and FU24 were included to investigate the change in performance from baseline to this follow-up. Baseline RBANS data was available for most of the participants for subsample A (N=90) and B (N=67). For the remaining three participants of subsample A and two participants of subsample B with missing baseline data, we used the RBANS data measured three months after baseline. Two participants in subsample A and one participant in subsample B were excluded from the investigation of cognition, as there were indications of possible mild cognitive impairment (MCI) between one and three years after baseline.

### 2.5. PET preprocessing and data preparation

Positron emission tomography (PET) imaging was conducted at the McConnell Brain Imaging Centre of the Montreal Neurological Institute (Quebec, Canada) utilizing a brain-dedicated Siemens PET/CT high-resolution research tomograph. Data acquisition and processing followed the methodology described by Yakoub and colleagues.^37^ Briefly, Aβ-PET images using 18F- NAV4694 (NAV) were acquired between 40 and 70 minutes post-injection, with an approximate injection dose of 6 mCi. Tau-PET images, utilizing 18F-flortaucipir (FTP), were captured 80 to 100 minutes post-injection, with an injection dose of around 10 mCi. The scans included 5-minute frames and an attenuation scan. PET images were reconstructed using a 3D ordinary Poisson ordered subset expectation maximum algorithm (OP-OSEM) with 10 iterations and 16 subsets, and were corrected for decay and motion. Scatter correction employed a 3D scatter estimation method. T1-weighted MRI images were segmented into 34 bilateral regions of interest (ROIs) based on the Desikan-Killiany atlas using FreeSurfer version 5.3. PET images were then realigned, temporally averaged, and co-registered to a T1-weighted image (specifically the scan closest in time to PET data acquisition). These images were then masked to exclude cerebrospinal fluid (CSF) signal and smoothed with a 6 mm Gaussian kernel. Standardized uptake value ratios (SUVRs) were calculated as the ratio of tracer uptake in the ROIs relative to the cerebellar gray matter for amyloid-PET scans, or to the inferior cerebellar gray matter for tau-PET scans. All PET data preprocessing was conducted using a standard pipeline available at https://github.com/villeneuvelab/vlpp. We assessed bilateral entorhinal FTP SUVR by averaging the uptake ratios of the left and right entorhinal cortices, and global NAV SUVR.

### 2.6. MRI data acquisition

MRI measurements were acquired with a Siemens Tim Trio 3-Tesla MRI scanner located at the Cerebral Imaging Centre of the Douglas Mental Health University Institute. A Siemens standard 12- or 32-channel coil was used (Siemens Medical Solutions, Erlangen, Germany). T1- weighted anatomical MPRAGE scans (TR = 2300 ms; TE = 2.98 ms; TI = 900 ms; 9° flip angle; FOV = 256×240×176 mm) were acquired with a 1 mm isotropic voxel resolution. Resting-state data were obtained using an EPI sequence (TR = 2000 ms; TE = 30 ms; a = 90◦; FOV = 256×256 mm; 32 slices), resulting in a 4 mm isotropic voxel resolution. The resting-state scan had a duration of 10 minutes. Additional fieldmaps were acquired to correct for spatial distortions of the functional data due to field inhomogeneities in the unwarping process.

### 2.7. MRI preprocessing

Functional and structural data were preprocessed using MATLAB and Statistical Parametric Mapping, version 12 (SPM12) (Functional Imaging Laboratory, UCL) as well as the CONN toolbox, version 22.a.^38^ The fMRI data were realigned to the first volume of the first session. Voxel-displacement maps were created using fieldmaps from the respective scanning session to correct susceptibility artifacts via unwarping. The functional data were co-registered to the structural T1-weighted images of the respective session. The structural images were segmented into different tissue types (gray matter, white matter, cerebrospinal fluid) and normalized to the Montreal Neurological Institute (MNI) reference frame for the human brain using the IXI549Space template implemented in SPM12 in the unified segmentation and normalization procedure of SPM12. The translation parameters derived from the structural images were then applied on the functional data for normalization to MNI space. Potential outliers due to excessive head motion were detected using ART, where volumes with framewise displacement above 0.5 mm/ TR or global intensity z score of 3 were flagged. Functional data were denoised after an anatomical component-based noise correction procedure (CompCor). Potentially confounding effects were regressed out using motion parameters and their first order derivatives, flagged outlier volumes, cerebral white matter, and areas containing cerebrospinal fluid. A band-pass filter of 0.008 Hz to 0.09 Hz was applied to minimize noise from physiological and motion sources.

### 2.8. First-level analyses and regions of interest

Connectivity matrices were derived for each individual participant at each session. Fisher z-transformed correlation coefficients for each pair of ROI (region of interest) time series were computed by ROI-to-ROI bivariate correlations and these formed the first-level matrices. This way, the strength of association (joint and opposing activation) between two ROIs can be investigated per participant, however, the associations are undirected. Convolution of the equal weights of the scans with a canonical hemodynamic response function (hrf-weighting) was applied to weight down the influence of the first volumes. ROIs for ROI-to-ROI functional connectivity analysis were selected a-priori based on a literature research on episodic memory brain areas. They include i) the MTL regions anterior and posterior hippocampus, lateral and medial parahippocampal cortex (PHC), entorhinal cortex (EC) and anterior and posterior perirhinal cortex (PRC); ii) the PMC regions precuneus, posterior cingulate cortex (PCC) and retrosplenial cortex (RSC) and iii) the medial prefrontal cortex (mPFC). The precuneus consisted of four subregions, the mPFC of three subregions. For a visualization of ROIs, see Figure 1B. The ROIs were derived from the Brainnetome atlas.^39^ As a control network, we used the visual network from the Yeo atlas.^40^ For atlas labels see Supplementary Table S1.

### 2.9. Second-level analyses: Assessment of the effects of aging and AD pathology

For each individual functional connection, a separate General Linear Model (GLM) was estimated. Our main questions regarding the MTL-PMC-prefrontal network were whether i) decreasing rsFC strength over time, especially with older age, is visible in the presence of low pathology (A^-^T^-^) and whether ii) increasing rsFC is associated with higher early AD pathology, especially in *APOE4* carriers.

In subsample A, we first investigated the i) FU12>baseline and ii) FU24>baseline contrast to identify connections with significant change in rsFC strength between sessions via paired t-tests in CONN. Second, we used change in rsFC over time as dependent variables in multiple linear regression models with baseline age as independent variable as well as the covariates *APOE* group, sex, education, and difference in days between baseline and the respective measurements. Third, we conducted an additional cross-sectional analysis of effects of baseline age on baseline rsFC accounting for *APOE* group, sex, and education.

Regarding the CSF-characterized subsample B, we used change in rsFC from baseline to FU24 as the dependent variable for multiple linear regression models. As independent variables of interest, we used i) change of pathology from baseline to FU24 and ii) pathology at baseline. We formed separate models to investigate the effect of Aβ_1-42_, p-tau_181_, and the p-tau_181_/Aβ_1-42_ ratio. Age, *APOE* group, sex, education, and difference in days between baseline and FU24 were included as covariates. Measures of change were calculated as difference scores, e.g. *pathology at FU24 - pathology at baseline*.

Inferential statistics were performed at the level of individual networks (groups of related connections) and based on the hypothesis defining the expected association. Network-level inferences were based on nonparametric statistics from Network Based Statistics analyses (NBS) which uses randomization of residuals of the second-level model to determine the significance of clusters of connections in CONN; 10000 iterations were used. We calculated the 95% confidence interval for the cluster *p*-value obtained from the permutation test (95% CI*_p_*) to quantify the uncertainty due to the finite number of permutations.^41^ Results were thresholded using a combination of a cluster-forming threshold of *p* < 0.001 at the connection level and a corrected threshold of *p*-FDR < 0.05 at the network level.^38^ For individual connections that were part of a significant cluster, computation of Cohen’s d (d) for t-tests and standardized β for regressions with the corresponding 95% confidence intervals (95% CI), assessment of the respective model requirements, and additional figure creation with the ggplot^42^ and ggseg^43^ packages were conducted using R (R Core Team) version 4.2.3, and RStudio, version 2022.07.1 (RStudio Team). The R code used for analyses is publicly available (https://github.com/fislarissa/MTL_PMC_longitudinal_rsFC). For linear models, we ensured that multicollinearity and heteroscedasticity were not present. Further, we tested for a normal distribution of residuals using the Shapiro-Wilk test on the standardized residuals of each model.

### 2.10. Assessment of graph measures

Additional graph theoretical measures were calculated with the CONN toolbox to assess changes in network properties over time. A graph consists of nodes and edges (connections) and represents the elements of a complex system and their interrelationships^44^. We focused on two main aspects: MTL-PMC-prefrontal network integration and subnetwork segregation (i.e. MTL, PMC and mPFC separately). For network integration, we calculated global efficiency, shortest path length, and nodal betweenness centrality. Global efficiency was measured to assess the efficiency of information transfer between nodes. Shortest path length was analyzed to determine the average minimum distance between nodes, with an increase indicating reduced network efficiency. Nodal betweenness centrality was used to identify how central specific nodes are to the network’s communication pathways. For subnetwork segregation, we focused on the clustering coefficient and local efficiency. The clustering coefficient was measured to evaluate the extent of local clustering within the network, indicating how well nodes form tightly-knit subgroups. Local efficiency was calculated to assess the efficiency of information transfer within these local subnetworks. To conduct the graph analysis, we specified in our preregistration^1^ to graphically determine the optimal primary threshold for each subsample (Achard and Bullmore, 2007) (see Supplementary Figure S1 for details). In short, we calculated global and local efficiency across cost thresholds for each subsample, comparing the measures for the real data to random and lattice networks. We used the formula *Global efficiency (Data) - Global efficiency (Lattice) + Local efficiency (Data) - Local efficiency (Random)*. The highest value, representing the greatest divergence between the real network and the modeled structures, was selected as the optimal cost threshold. This resulted in a cost threshold of 0.14 for subsample A and 0.16 for subsample B, which were then applied to the respective second-level rsFC matrix to establish the adjacency matrix. The cost threshold means that the 14% or 16% largest correlation coefficient values were used to build the binary adjacency matrix, which either uses 0 or 1 to define whether there is a suprathreshold connection between the respective pair of nodes. The same models as described for the rsFC analysis were tested and again based on the hypothesis defining the expected association. A second-level analysis threshold of *p* < 0.05 and FDR-correction for multiple univariate testing was applied.

### 2.11. Assessment of cognitive performance

We investigated the association of cognitive performance over time with rsFC strength (i) at baseline and (ii) over time. Our two remaining hypotheses were centered around the association of longitudinal memory performance and rsFC strength. These were as follows: iv) mild age-related (A^-^T^-^) decline or less practice effects in episodic memory performance are associated with steeper decrease in rsFC strength over time, especially in older age, and v) higher or increasing rsFC strength could be an initial beneficial or compensatory process if predicting maintained episodic memory performance or a detrimental process if predicting decline in performance. All statistical analyses regarding cognition were conducted using RStudio. The R code used for analyses is publicly available (https://github.com/fislarissa/MTL_PMC_longitudinal_rsFC). For linear models, we ensured that multicollinearity and heteroscedasticity were not present. Further, we tested for a normal distribution of residuals using the Shapiro-Wilk test on the standardized residuals of each model. We used a paired t-test in each subsample to analyze change in memory performance over two years. Then we extracted the first-level rsFC matrices from CONN and used (i) the functional connection that showed the strongest effect of aging over two years in subsample A and (ii) the functional connection that showed the strongest effect for AD pathology over two years in subsample B. These connections were RSC - PCC and anterior hippocampus - superior precuneus, respectively. We implemented this approach to avoid multiple testing.

First, addressing hypothesis v) we used multiple regression to predict change in the RBANS delayed memory index score over two years, from baseline to FU24, with the respective measure of rsFC strength at baseline, taking age, *APOE*, sex, education, time from baseline to FU24, and a potential interaction of FC\**APOE* into account. For subsample B, we included the p-tau_181_/Aβ_1-42_ ratio as an additional covariate. Second, addressing hypotheses iv) and v), we predicted change in RBANS performance over two years with change in rsFC strength over two years using the covariates age, sex, education, and time from baseline to FU24. Then, we repeated these models while including the AD-related covariates *APOE* and the interaction terms FC\**APOE* and FC*age in subsample A and *APOE*, the p-tau_181_/Aβ_1-42_ ratio, FC\**APOE* and FC*p-tau_181_/Aβ_1- 42_ ratio in subsample B. Third, as a control analysis, we used the change in attention index score from baseline to FU24 as outcome variable in the same models as described for the delayed memory index. Measures of change were calculated as difference scores, e.g. *RBANS at FU24 - RBANS at baseline*.

### 2.12. Assessment of structural changes of the hippocampus

As a control analysis we investigated structural change in the whole bilateral hippocampus as a proxy for early neurodegenerative processes. We utilized FreeSurfer 7.1 (Laboratory for Computational Neuroimaging, Athinoula A. Martinos Center) and its longitudinal pipeline^45^ to analyze changes in hippocampus volume across the three sessions (baseline, FU12, FU24). For one subject from subsample B, there was no MRI data for FU12 available. Initially, T1-weighted MRI scans for each session were processed using the standard cross-sectional stream to generate anatomical reconstructions. An unbiased within-subject template was then created from the images of all three sessions, serving as a reference to align and refine segmentations. This approach minimizes inter-scan variability and enhances the accuracy of longitudinal measurements. Hippocampal volumes were extracted from the segmented images for each session, registered to the within-subject template. For subsample A and B separately, we used a rmANOVA to test for changes in hippocampus volume over the three sessions. We then performed post hoc t-tests for a significant ANOVA. To assess the normalized rate of change of hippocampus volume over two years, we used the formula *(volume at FU24 - volume at BL)/ volume at BL*100*. We then (i) correlated rate of change of hippocampus volume with AD pathology, (ii) used rate of change of hippocampus volume instead of change in rsFC in the models described above and (iii) included rate of change of hippocampus volume as a covariate in the models predicting change in rsFC or memory performance.

## 3. Results

### 3.1. Assessment of the effects of aging on rsFC

#### 3.1.1. Assessment of change in rsFC over one and two years

In the AD-biomarker negative subsample A, rsFC decreased significantly in two clusters, both over one year (from baseline to FU12) and over two years (from baseline to FU24). The first cluster showed rsFC decrease between superior mPFC - PMC and within PMC, the second cluster showed rsFC decrease between MTL - precuneus.

For baseline to FU12, cluster 1 (*p* = 0.008, [95%CI*_p_* 0.006, 0.01], see Figure 2A) consisted of the left superior mPFC and three regions of the PMC. Specifically, rsFC decreased between left superior mPFC - right PCC, left superior mPFC - left PCC, and left superior mPFC - left RSC (see Table 2 for statistics). Cluster 2 (p = 0.008, [95%CI*_p_* 0.006, 0.01], see Figure 2B) consisted of the bilateral lateral PHC and the medial precuneus. Specifically, rsFC decreased between left lateral PHC - left medial precuneus, left lateral PHC - right medial precuneus, and right lateral PHC - left medial precuneus (see Table 2 for statistics).

**Fig 2.**
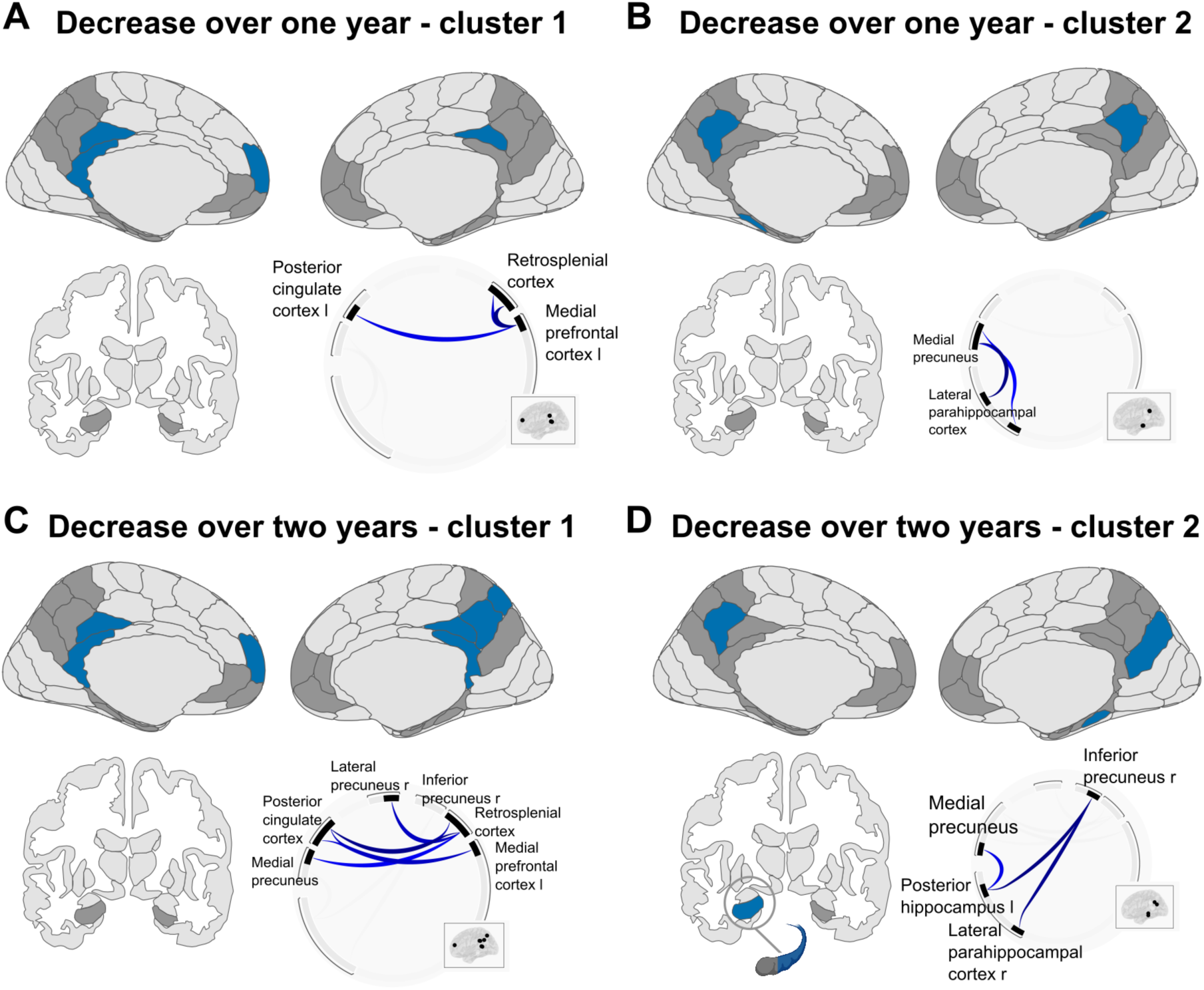
Longitudinal decrease in resting-state functional connectivity in amyloid- and tau-negative older adults. Illustrations are presented in neurological view. **A)** Cluster 1 for decrease over one year, from baseline to the follow-up after 12 months. **B)** Cluster 2 for decrease over one year, from baseline to the follow-up after 12 months. **C)** Cluster 1 for decrease over two years, from baseline to the follow-up after 24 months. D) Cluster 2 for decrease over two years, from baseline to the follow-up after 24 months. l = left. r = right. See Table 2 for statistics.

**Table 2.**
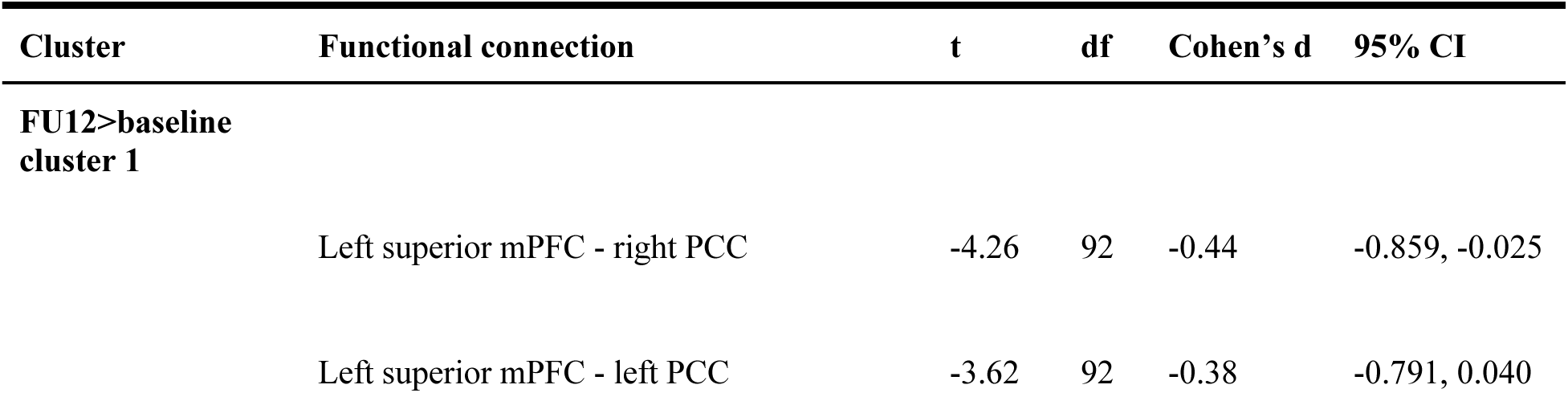
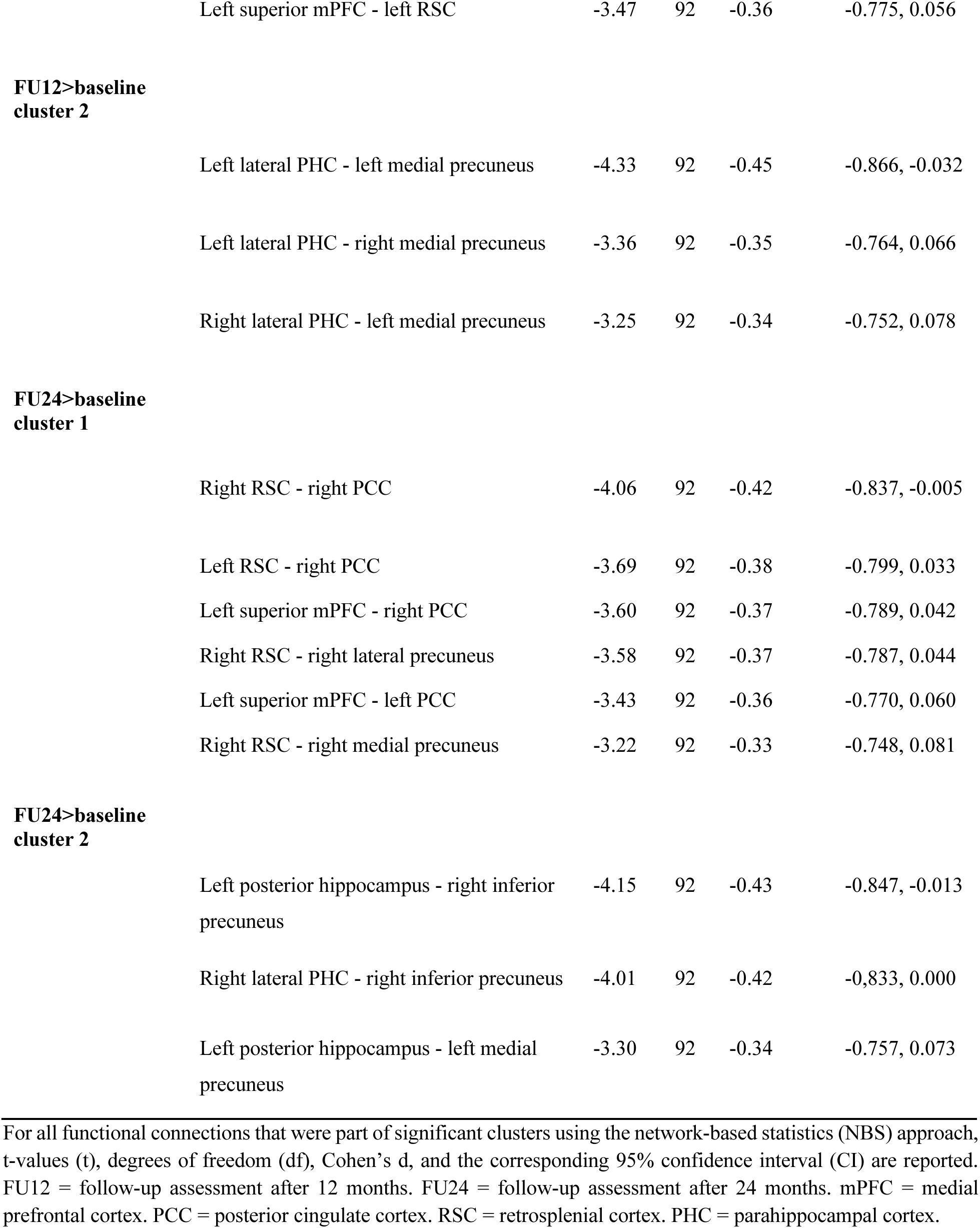
Statistical reporting for longitudinal decrease in functional connectivity with aging.

Regarding the difference in rsFC strength from baseline to FU24, there were again two significant clusters showing decrease in FC. Cluster 1 (*p* = 0.001, [95%CI*_p_* 0.001, 0.002], see Figure 2C) consisted of ROIs within the PMC and the superior mPFC. The strongest decrease of rsFC was observed between right RSC - right PCC (see Table 2 for the other connections). Cluster 2 (*p* = 0.005, [95%CI*_p_* 0.003, 0.006], see Figure 2D) consisted of the posterior MTL and the inferomedial precuneus. Specifically, rsFC decreased between left posterior hippocampus - right inferior precuneus, right lateral PHC - right inferior precuneus, and left posterior hippocampus - left medial precuneus (see Table 2 for statistics).

In summary, we observed decreasing rsFC strength over one and two years in AD- biomarker negative older adults. This included decreasing rsFC between the mPFC and the PMC, between the MTL and the PMC and within the PMC subnetwork.

Within our control network, the visual network, there were no significant changes in rsFC over time (p > 0.05).

#### 3.1.2. Assessment of the effects of baseline age on change in rsFC over one and two years

There was a significant effect of baseline age on change in rsFC over one year, from baseline to FU12, for a cluster including five ROIs (*p* = 0.034, [95%CI*_p_* 0.030, 0.038], see Figure 3A and Tables S2-4), with involvement of the PHC in every connection. In this cluster, older age was related to a steeper decrease in rsFC over time. The specific connections were right lateral PHC - left posterior PRC (β= −0.34, t(87) = −3.15, [95%CI −0.553, −0.125], see Figure 3B), right medial PHC - right PCC (β=−0.32, t(87) = −3.03, [95%CI −0.537, −0.112]), and right lateral PHC - left PCC (β= −0.33, t(87) = −3.03, [95%CI −0.541, −0.112]). There were no significant effects of baseline age on change in rsFC over two years from baseline to FU24. Further, there were no significant effects of *APOE* group, sex, or education neither for change in rsFC from baseline to FU12 nor for baseline to FU24. In an additional analysis, we did not find significant differences in baseline rsFC due to age (*p* > 0.05). In our control network, the visual network, there were no significant effects of age on change in rsFC (*p* > 0.05).

**Figure 3.**
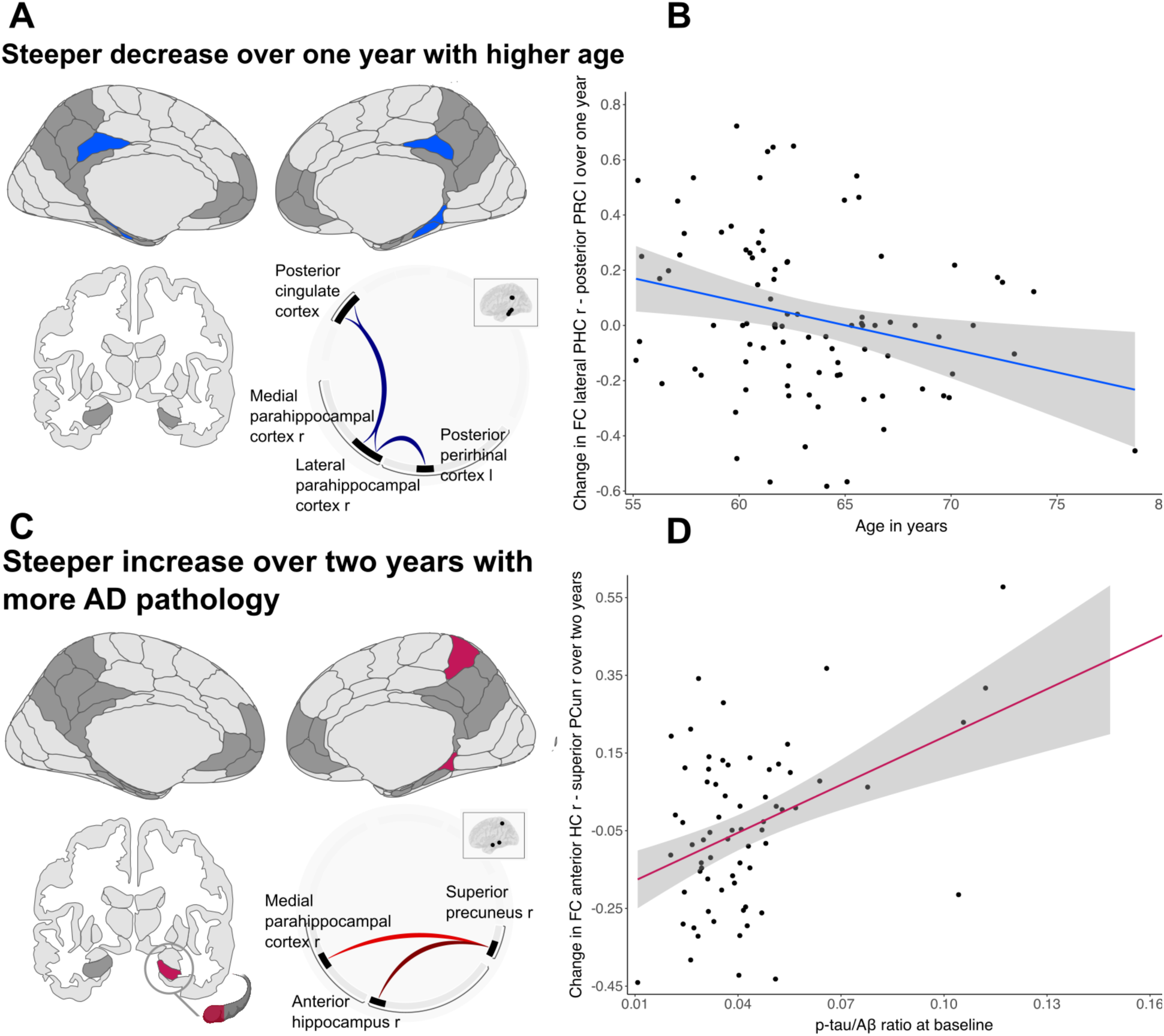
Longitudinal change in functional connectivity with older age and Alzheimer’s pathology. Illustrations are presented in neurological view. **A)** In the amyloid- and tau-negative older adults of subsample A, there was a steeper longitudinal decrease over one year in resting-state functional connectivity (rsFC) with higher age at baseline. Involved brain areas are depicted in blue. **B)** Change in rsFC over one year was negatively associated with age at baseline. The connection showing the strongest effect (lateral parahippocampal cortex - posterior perirhinal cortex) is used for visualization. Residuals of the model were normally distributed, multicollinearity and heteroscedasticity were not present. **C)** In the amyloid and tau cerebrospinal fluid (CSF) characterized older adults of subsample B, there was a steeper longitudinal increase over two years in rsFC with a higher Alzheimer’s pathology (AD) ratio of p-tau/amyloid measured in CSF. Involved brain areas are depicted in red. **D)** Change in rsFC over two years was positively associated with the p-tau/amyloid ratio at baseline. The connection showing the strongest effect (anterior hippocampus - superior precuneus) is used for visualization. Residuals of the model were normally distributed, multicollinearity and heteroscedasticity were not present. l = left. r = right. FC = functional connectivity. PHC = parahippocampal cortex. PRC = perirhinal cortex. AD = Alzheimer’s disease. Aβ = amyloid-beta. HC = hippocampus. PCun = precuneus.

### 3.2. Assessment of the effects of AD pathology on change in rsFC over two years

In the CSF-characterized subsample B, there was no association of change in rsFC strength with change in Aβ_1-42_, p-tau_181_ or the p-tau_181_/Aβ_1-42_ ratio (all *p* > 0.05). For all variables, we investigated change over two years, from baseline to FU24. Importantly, all three pathology measures themselves did not change significantly over this timespan (all *p* > 0.05). We therefore investigated whether the respective measure of pathology at baseline showed an effect on change in rsFC over two years. We observed no apparent effect of baseline Aβ_1-42_ on change in rsFC (*p*>0.05). For baseline p-tau_181_ however, the connection between right anterior hippocampus - right superior precuneus showed the strongest effect that survived the connection threshold of *p* < 0.001 (β= 0.44, t(62) = 3.81, [95%CI 0.208, 0.666]) but not the cluster threshold (*p* = 0.089, [95%CI*_p_* 0.083, 0.095]). For the p-tau_181_/Aβ_1-42_ ratio, a cluster covering three ROIs of the MTL-PMC network was significant (*p* = 0.032, [95%CI*_p_* 0.029, 0.035], see Figure 3C and Tables S5-6). In this cluster, a higher p-tau_181_/Aβ_1-42_ ratio was related to increasing rsFC strength over two years between right anterior hippocampus - right superior precuneus (β=0.51, t(62) = 4.190, [95%CI 0.269, 0.759]), as depicted in Figure 3D, and between right medial PHC - right superior precuneus (β= 0.44, t(62) = 3.371, [95%CI 0.178, 0.696]).

In our control network, the visual network, there was no significant effect of AD pathology on change in rsFC (all *p* > 0.05).

### 3.3. Assessment of change in graph measures over one and two years

The only graph measure that showed significant longitudinal changes over time was global efficiency, a measure of integration of nodes within a network. Global efficiency is the normalized average inverse shortest path-length of a node with each of the other nodes of the network. There were decreases in global efficiency of the network for subsample A over one year, from baseline to FU12 (t(92) = −1.86, *p* = 0.033), as well as marginally significant over two years, from baseline to FU24 (t(92) = −1.65, *p* = 0.051). From baseline to FU12, the nodes significantly contributing to the decrease in global efficiency were the bilateral superior mPFC and the left anterior PRC (see Figure 4A). From baseline to FU24, the nodes significantly contributing to the decrease were the RSC, superior mPFC, PCC and medial precuneus, all bilateral (see Figure 4B). Further, there was an effect of age on global efficiency of the network from baseline to FU12 (t(87) = −2.21, *p* = 0.02) with the bilateral posterior PRC and the bilateral EC (see Figure 4C) significantly contributing, such that a stronger decrease in global efficiency was present in older participants. Statistical reporting for each node can be found in Table 3. There was no significant age effect for change from baseline to FU24 and no effect of *APOE4* group on change in graph measures. For subsample B, there was no effect of AD pathology or *APOE4* group on change in graph measures over two years. We note that the relatively small network size (16 nodes) may limit the generalizability of our graph theoretical results, and care should be taken when interpreting these findings in the context of larger or more complex brain networks.

**Figure 4.**
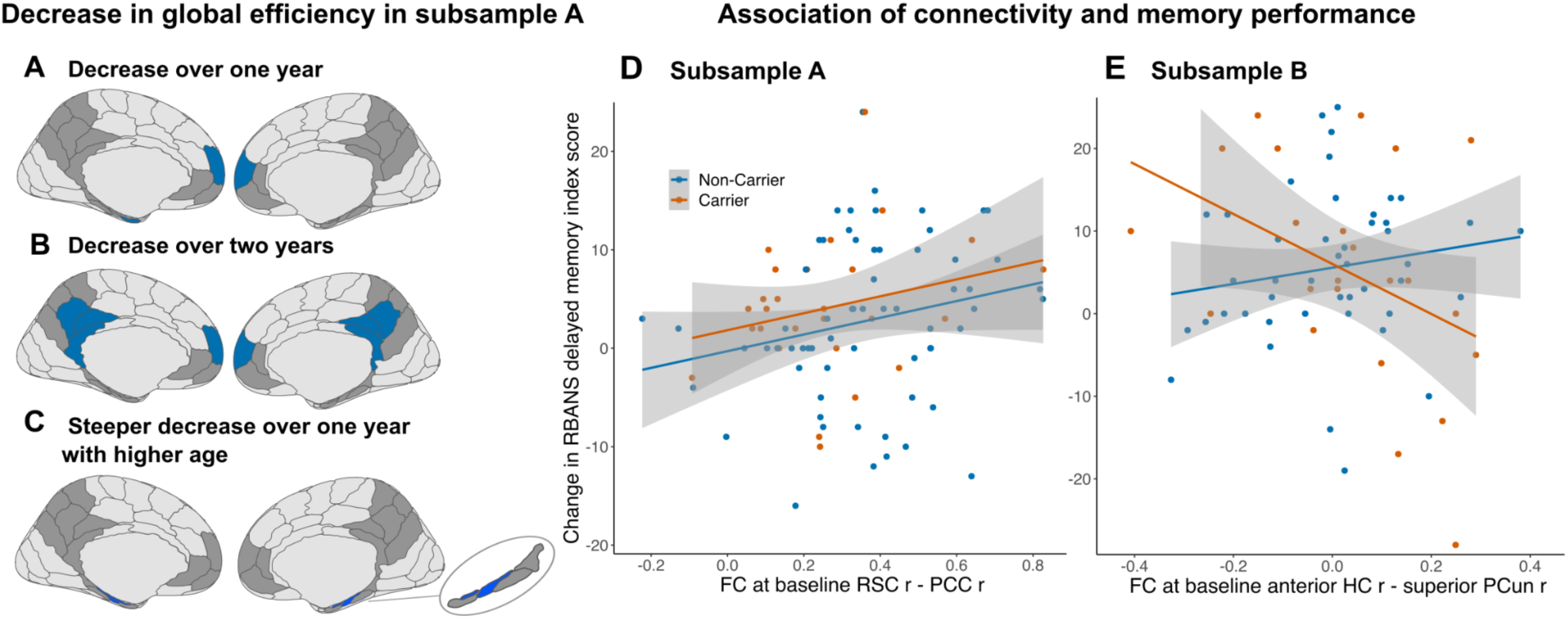
Longitudinal decrease in global efficiency in amyloid and tau negative older adults and association of functional connectivity and change in episodic memory performance. Illustrations are presented in neurological view. As there was no involvement of the hippocampus, the coronal view is not shown. **A)** Decrease over one year, from baseline to the follow-up after 12 months, in the left and right superior mPFC and left anterior PRC. **B)** Decrease over two years, from baseline to the follow-up after 24 months, in the RSC, superior mPFC, PCC, and medial PCun, each in the left and right hemisphere. **C)** Steeper longitudinal decrease over one year with higher age at baseline for the posterior PRC and EC, each in the left and right hemisphere. **D)** In the amyloid- and tau-negative older adults of subsample A, there was an association of functional connectivity (FC) at baseline and change in episodic memory performance measured with the RBANS. There was no effect of *APOE* genotype group. The functional connection with the strongest decrease over time, right RSC - right PCC, was investigated. N=91. **E)** In the amyloid and tau cerebrospinal fluid (CSF) characterized older adults of subsample B, there was an interaction of resting-state functional connectivity at baseline and *APOE* group on change in episodic memory performance. N=86. mPFC = medial prefrontal cortex. PRC = perirhinal cortex. RSC = retrosplenial cortex. PCC = posterior cingulate cortex. PCun = precuneus. EC = entorhinal cortex. RBANS = Repeatable Battery for Assessment of Neuropsychological Status. r = right. RSC = retrosplenial cortex. PCC = posterior cingulate cortex. HC = hippocampus.

**Table 3.**
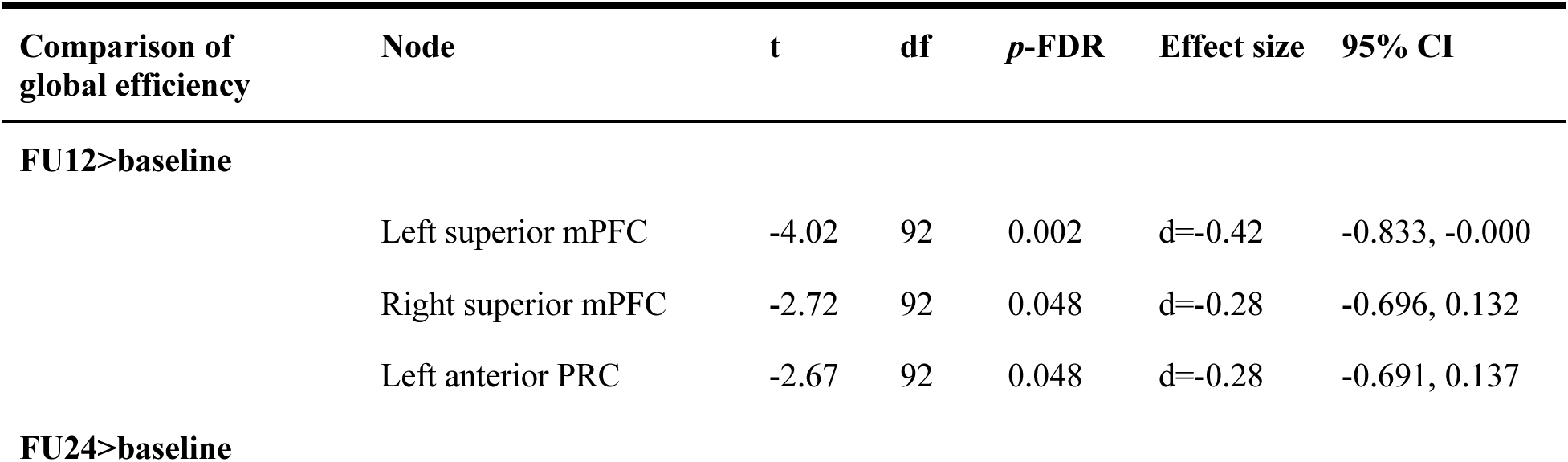
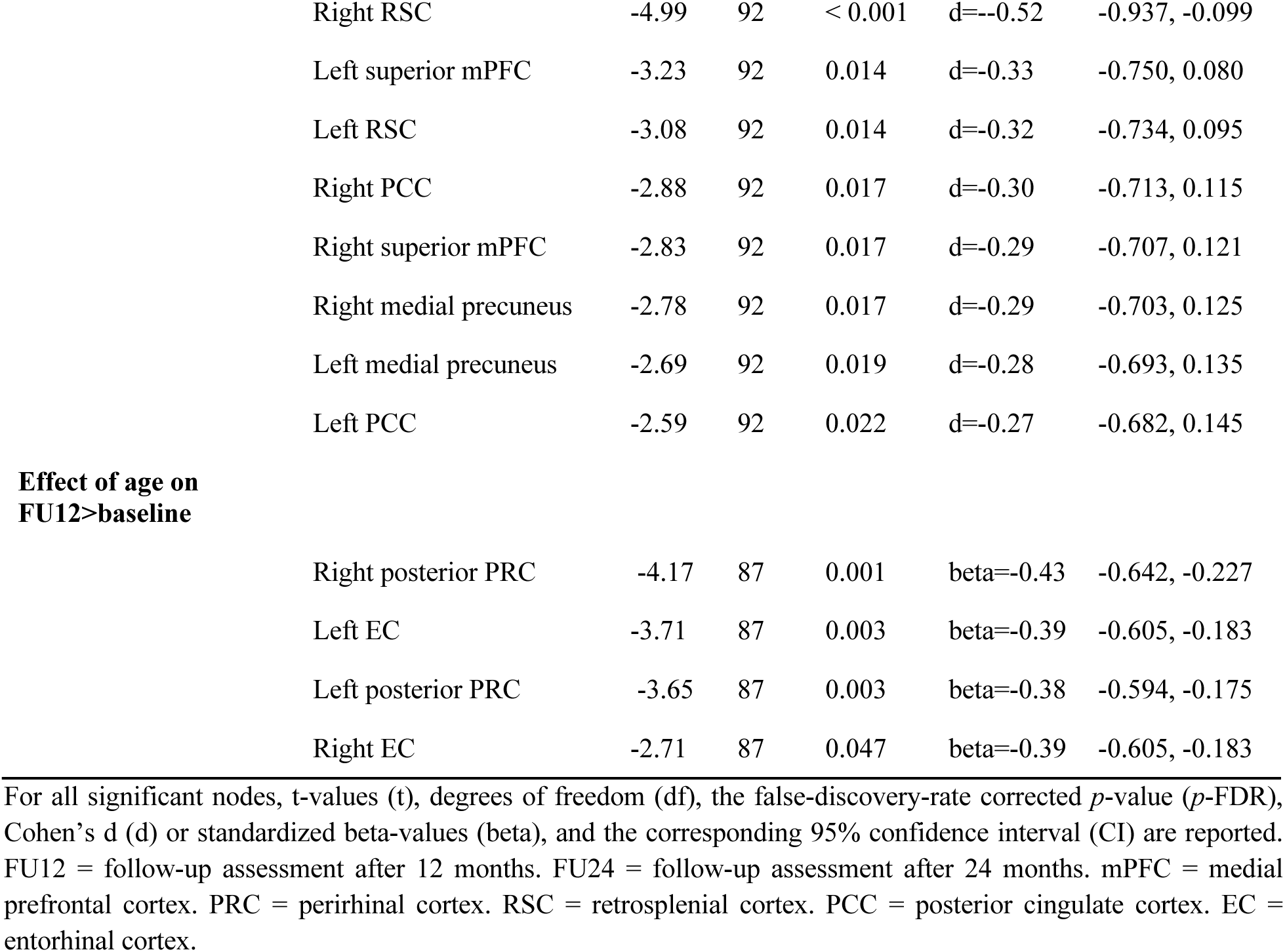
Statistical reporting for nodes contributing to longitudinal decrease in global efficiency with aging.

| Comparison of global efficiency | Node | t | df | p-FDR | Effect size | 95% CI |
| --- | --- | --- | --- | --- | --- | --- |
| **FU12>baseline** |  |  |  |  |  |  |
|  | Left superior mPFC | -4.02 | 92 | 0.002 | d=-0.42 | -0.833, -0.000 |
|  | Right superior mPFC | -2.72 | 92 | 0.048 | d=-0.28 | -0.696, 0.132 |
|  | Left anterior PRC | -2.67 | 92 | 0.048 | d=-0.28 | -0.691, 0.137 |
| **FU24>baseline** |  |  |  |  |  |  |

|  |  |  |  |  |  |
| --- | --- | --- | --- | --- | --- |
| Right RSC | -4.99 | 92 | < 0.001 | d=-0.52 | -0.937, -0.099 |
| Left superior mPFC | -3.23 | 92 | 0.014 | d=-0.33 | -0.750, 0.080 |
| Left RSC | -3.08 | 92 | 0.014 | d=-0.32 | -0.734, 0.095 |
| Right PCC | -2.88 | 92 | 0.017 | d=-0.30 | -0.713, 0.115 |
| Right superior mPFC | -2.83 | 92 | 0.017 | d=-0.29 | -0.707, 0.121 |
| Right medial precuneus | -2.78 | 92 | 0.017 | d=-0.29 | -0.703, 0.125 |
| Left medial precuneus | -2.69 | 92 | 0.019 | d=-0.28 | -0.693, 0.135 |
| Left PCC | -2.59 | 92 | 0.022 | d=-0.27 | -0.682, 0.145 |
| **Effect of age on FU12>baseline** |  |  |  |  |  |
| Right posterior PRC | -4.17 | 87 | 0.001 | beta=-0.43 | -0.642, -0.227 |
| Left EC | -3.71 | 87 | 0.003 | beta=-0.39 | -0.605, -0.183 |
| Left posterior PRC | -3.65 | 87 | 0.003 | beta=-0.38 | -0.594, -0.175 |
| Right EC | -2.71 | 87 | 0.047 | beta=-0.39 | -0.605, -0.183 |

For all significant nodes, t-values (t), degrees of freedom (df), the false-discovery-rate corrected *p*-value (*p*-FDR), Cohen's d (d) or standardized beta-values (beta), and the corresponding 95% confidence interval (CI) are reported. FU12 = follow-up assessment after 12 months. FU24 = follow-up assessment after 24 months. mPFC = medial prefrontal cortex. PRC = perirhinal cortex. RSC = retrosplenial cortex. PCC = posterior cingulate cortex. EC = entorhinal cortex.

### 3.4. Assessment of change in cognitive performance over two years

There was a change in the mean memory performance in both subsamples over two years from baseline to FU24. Specifically, we observed a slight increase in performance in subsample A (t(90) = 3.84, *p* < 0.001, d=0.37, [95%CI 0.174, 0.569]) from a mean score of 104 (SD=9) to 107 (SD=8) and in subsample B (t(67) = 4.11, *p* < 0.001, d=0.59, [95%CI 0.280, 0.890]) from a score of 102 (SD=10) to 107 (SD=8).

When examining associations between change in memory performance and rsFC (at baseline and over time), we limited our analysis to the connection that showed the strongest decrease in rsFC with aging in subsample A (i.e. RSC - PCC), and the connection that showed the strongest relationship to AD pathology in subsample B (i.e. anterior hippocampus - superior precuneus).

With respect to rsFC at baseline, a steeper increase in memory performance was predicted by higher RSC - PCC rsFC in subsample A (β=0.24, t(84)=2.15, *p*=0.035, [95%CI 0.018, 0.455], see Figure 4D and supplementary Table S7). No interaction between *APOE* group and baseline rsFC on memory change was observed in subsample A for this connection. For subsample B, baseline rsFC between anterior hippocampus - superior precuneus was not related to performance change in the whole sample (*p*>0.05). However, we found an interaction of rsFC and *APOE* group on change in memory performance (β=-0.65, t(59)=-2.54, *p*=0.014, [95%CI −1.169, −0.139], see Figure 4E and supplementary Table S8). Specifically, in *APOE4* carriers, higher baseline rsFC tended to be related to less increase in performance, although the effect was only marginal (β=- 0.52, t(16)=-2.08, *p*=0.054, [95%CI −1.058, 0.011], see Figure 4E orange line and supplementary Table S9). As the subgroup of 23 *APOE4* carriers was small, we used additional bootstrapping with 1000 replications to generate 95%CIs of the estimate [95%CI_b_ −114.268, 4.482]) (see supplementary Figure S2). In non-carriers (N=65), there was no effect of baseline rsFC on change in memory performance (*p*>0.05; Fig. 4E blue line). Regarding AD pathology, the p-tau_181_/Aβ_1-42_ ratio at baseline was not significantly related to change in performance (*p*>0.05).

With respect to change in rsFC over time, there was no significant association with change in episodic memory performance for subsample A or B when only accounting for age, sex, education, and time from baseline to FU24. Furthermore, there was no significant effect of change in rsFC over time on change in memory performance when including the AD-related covariate *APOE4* in subsample A and both *APOE4* and the p-tau_181_/Aβ_1-42_ ratio in subsample B. Additionally, there were no interaction effects regarding change in rsFC strength predicting change in memory of FC\**APOE* or FC*age in subsample A and FC\**APOE* or FC*p-tau_181_/Aβ_1-42_ ratio in subsample B (all *p*>0.05, see supplementary Tables S10-13).

Lastly, we conducted the reported analyses with the RBANS attention index score instead of the delayed memory index score. We found no change in the attention score over time and no effects of rsFC at baseline, rsFC change or interactions of rsFC and *APOE* group on change in the attention score (all *p*>0.05, see supplementary Tables S14-19).

### 3.5. Assessment of structural changes of the hippocampus over two years

Finally, we examined whether the associations of aging, AD pathology and memory performance with rsFC strength were influenced by early structural neurodegeneration as a control analysis. Using whole bilateral hippocampus volume as a proxy of early neurodegenerative processes, we found a significant reduction thereof over two years in subsample A (F(2,184)=27.93, *p*<0.001, η_g_^2^=0.0022) and subsample B (F(2,134)=12.51, *p*<0.001, η_g_^2^=0.0012). Bonferroni-corrected post-hoc t-tests revealed significant reductions in hippocampus volume between all sessions, except for subsample B from baseline to FU12 (see supplementary Table S20). In subsample B, the rate of change of hippocampus volume was correlated with the p-tau_181_/Aβ_1-42_ ratio at baseline (r=-0.25, t(67)=-2.11, *p*=0.038, [95%CI −0.46, −0.02]), but not with p-tau or amyloid on their own (*p*>0.05). When including age, *APOE* group, sex, education, and time between baseline and FU24 in a multiple regression predicting the rate of change of hippocampus volume, the only significant predictor of rate of change was time between the two measurements (β=-0.34, t(62)=-2.82, *p*=0.007, [95%CI −0.57, −0.10], see supplementary Table S21). Although a reduction in hippocampus volume was observed, the rate of decline of hippocampus volume was not associated with change in rsFC or episodic memory performance when included as a covariate in the linear models described above (all *p*>0.05). Details can be found in supplementary Tables S22 - S31.

## 4. Discussion

### 4.1. Summary

In our longitudinal study assessing MTL-PMC-prefrontal rsFC strength and episodic memory performance, we observed a decrease in rsFC strength over one and two years related to aging in the absence of AD pathology. This decrease was regionally specific to FC within PMC and between posterior hippocampus - inferomedial precuneus. Furthermore, we found specific medial PHC - superior precuneus and anterior hippocampus - superior precuneus increase in rsFC strength over two years related to higher AD pathology. Regarding episodic memory performance over time, higher rsFC at baseline within the PMC was beneficial in older adults without evidence of AD pathology. However, our findings in the subsample with AD biomarkers indicated that higher rsFC between MTL and PMC might be detrimental for *APOE4* carriers, but not for *APOE4* non-carriers. Our results were specific to the memory network and memory performance, as there were no changes in our control network, the visual network, and no relations to performance in an independent attention test. We observed no significant effect of longitudinal hippocampal volume changes on these associations.

### 4.2. Effects of aging and baseline age on functional connectivity

Aging independent of AD pathology was related to longitudinal decrease in FC. Hypoconnectivity mainly occured between posterior MTL and inferomedial precuneus, within PMC and between superior mPFC and PCC. Consistent with our longitudinal results, previous cross-sectional studies found lower rsFC between hippocampal and neocortical regions in older age ^46^. More specifically, in accordance with our findings, recent studies reported that older age is related to lower rsFC within PMC and MTL, and between posterior MTL and PMC in amyloid-negative cognitively normal older adults, with the PHC as part of the PM system being particularly vulnerable to aging effects.^27,28^ Consistently, in our sample, PHC-PRC and PHC-PMC rsFC strength decreased more strongly over time in older individuals. Furthermore, for two nodes within the MTL, i.e. PRC and EC, we observed a steeper decrease in global efficiency with higher age, indicating network disconnectivity and inefficient exchange of information. This suggests a greater age-related vulnerability of the parahippocampal gyral regions to disintegration from the episodic memory network in the context of “normal” aging.^47^ The PHC is a major hub that functionally connects the MTL with PMC regions^48^ and seems to be a key region for age-related steeper decline in FC. In a previous study, lower rsFC and network dedifferentiation have been associated with higher age in the whole PM network cross-sectionally.^49^ Network dedifferentiation is associated with cognitive decline and refers to less distinctive neural representations.^50–52^ Similarly, we observed decreased global efficiency in PMC nodes, consistent with prior aging graph analysis.^53^

### 4.3. Effect of Alzheimer’s pathology on functional connectivity

While we observed little change in AD pathology measured via CSF over two years, we do report a significant increase in rsFC over the same period. This suggests that the specific observed MTL-PMC hyperconnectivity may be driven by existing pathological burden. Consistent with the hypothesis of a vicious cycle of pathology accumulation, aberrant activity and FC,^3,20,30^ we recently found that higher and increasing precuneus activity during memory retrieval was linked to subsequent higher Aβ burden, indicating that increased activity might contribute to Aβ accumulation.^54^ In the present study, increasing rsFC strength of the anterior hippocampus and of the PHC with the superior precuneus could be related to higher hyperphosphorylated tau in relation to Aβ levels. Soluble CSF p-tau_181_ serves as a marker of abnormal tau phosphorylation, a process that precedes the formation of neurofibrillary tangles and is closely associated with Aβ plaques.^55,56^ Recent findings suggest that CSF p-tau may drive the accumulation of tau tangles related to CSF Aβ pathology.^57^ Hyperphosphorylated tau is produced and secreted by neurons that, while still functional, are already compromised by Aβ accumulation and neuronal hyperexcitation.^58,59^ CSF Aβ_1-42_ is a marker for soluble Aβ peptides that are prone to aggregation into plaques.^60^

Interestingly, we observed subregion-specific rsFC changes between MTL and precuneus depending on the involvement of AD pathology (versus non-pathological aging). Specifically, MTL-superior precuneus rsFC increase was related to higher AD pathology, whereas MTL- inferomedial precuneus rsFC decrease was related to aging. Not much is known about functional subregions of the precuneus. Cross-sectionally, a previous study in the PREVENT-AD cohort reported a superior-inferior FC gradient of PMC to cortex, where changes in FC involving superior PMC were specifically related to tau burden.^61^

Specifically for the MTL-PMC network, a model of dysconnectivity and decreasing MTL- PMC FC strength with early amyloid in PMC was proposed previously,^62^ leading to local hyperactivation and aberrant FC within MTL due to the lack of neocortical inhibition, which drives tau spread from the MTL to functionally connected regions. This model is, however, based on cross-sectional data. Early tau seems to spread between highly functionally connected regions,^16,18,57^ contributing to a vicious cycle of pathology accumulation, later cell death and hypoconnectivity and -activity in advanced disease stages.^30^ Dynamic causal modeling (DCM) during an fMRI memory task showed reduced inhibition from PMC to MTL regions with increasing amyloid burden, this directed “hyperexcitation” in turn predicted MTL tau accumulation.^63^

### 4.4. Effects of functional connectivity and *APOE* genotype on change in cognition

We observed a change in memory performance over two years in both subsamples with a slight increase in performance, possibly reflecting practise effects.^64^ Higher rsFC strength within PMC at baseline seemed to be beneficial for longitudinal cognition regardless of *APOE4* genotype in older adults without AD pathology. This is consistent with previous cross-sectional and longitudinal findings, which have shown that memory-task activity and rsFC in older adults that more closely resemble activity and rsFC of younger adults are beneficial.^17,65–67^ In younger adults, however, no associations were found between cognitive performance and FC in a previous study, suggesting that FC strength may play a more significant role in older age.^68^

We further observed differential relationships on cognition for rsFC between MTL and precuneus depending on *APOE* genotype. Higher rsFC strength between anterior hippocampus and superior precuneus at baseline tended to be detrimental in *APOE4* carriers, but not in *APOE4* non-carriers. We also observed more AD pathology in *APOE4* carriers, which was, however, itself not related to memory performance. Furthermore, there were no significant differences in FC or cognition between *APOE* groups, as reported before.^69,70^ The differing FC-behavior relationship we observed could be due to subtle neurobiological changes related to early AD-pathology in *APOE4* carriers. Previous cross-sectional findings reported mixed results regarding the relationship between MTL-PMC rsFC and memory performance in dependence of *APOE4* genotype. In some studies, better memory performance was associated with higher hippocampus-PMC rsFC in older age regardless of *APOE* genotype.^71,72^ On the contrary, better episodic memory performance was related to lower rsFC of the temporal default mode network (DMN), only in *APOE4* carriers.^73^ We did not observe an effect of change in rsFC on change in memory performance over time, and no interaction of change in rsFC with *APOE*, age or AD pathology. In summary, our findings do not support the hypothesis that greater FC between anterior hippocampus and superior precuneus is beneficial or compensatory in the presence of AD risk as it was not related positively but rather negatively to future change in memory performance in *APOE4* carriers.^74^ However, the effect of change in FC on trajectories of cognition should be further investigated in long-term studies.

### 4.5. Limitations

First, we acknowledge that when investigating resting-state fMRI, there is limited information on cognitive processes during scanning. However, rsFC can provide valuable insights into brain dynamics relevant for cognition^70^ and functional changes have been reported in cognitively normal older adults both during cognitive tasks^75^ and at rest^70^. Future studies could directly extend our findings by assessing FC during memory-task performance and relate these findings to longitudinal amyloid, tau, behavior, and *APOE* genotype.

Second, our sample size was limited and predefined as we used an existing dataset. However, we preregistered our a-priori hypotheses and methods to strengthen our study. Further research could focus on more fine-grained regional differences and investigate hippocampal and PMC subregions.

Third, the PREVENT-AD cohort consists of primarily white, highly educated female participants, thus representing only a subsample of the general population. Future research should aim to replicate our findings in more ethnically and sociodemographically diverse cohorts.

Fourth, our subsample of A^-^T^-^ participants show variation in their sub-threshold levels of amyloid and tau, ranging from no detectable pathology to slightly below cut-off. Despite this limitation, our study contributes much needed longitudinal findings, as AD markers and especially tau markers were often not available in previous studies.

### 4.6. Conclusion

In summary, our results highlight region-specific differences in longitudinal rsFC strength between “normal” aging and early AD pathology. A decrease in within-PMC rsFC occurs with aging, a weaker decrease appears to be beneficial for episodic memory performance. The parahippocampal gyrus seems to be especially affected by decrease in rsFC in older age. Higher AD pathology is related to increase in MTL-PMC rsFC, specifically anterior hippocampus-superior precuneus FC. These findings shed light on the earliest changes in rsFC strength with AD pathology, which might already be disadvantageous for episodic memory performance in *APOE4* carriers.

## Acknowledgements

We want to thank the participants of the PREVENT-AD study for their time and effort as well as the researchers involved in building up the cohort https://preventad.loris.ca/acknowledgements/acknowledgements.php?DR=7.0.

## Data availability statement

Some of the data used in this manuscript is openly available at https://openpreventad.loris.ca. The remaining data can be shared upon approval by the scientific committee at the Centre for Studies on Prevention of Alzheimer’s Disease (StoP-AD) at the Douglas Mental Health University Institute after registration at https://registeredpreventad.loris.ca. Code for PET preprocessing is available at https://github.com/villeneuvelab/vlpp and for statistical analysis is avalable at https://github.com/fislarissa/MTL_PMC_longitudinal_rsFC.

