## Supplementary Material for "Differential effects of aging, Alzheimer’s pathology, and *APOE4* on longitudinal functional connectivity and episodic memory in older adults"

-

### Supplementary

Larissa Fischer[1], Jenna N. Adams[2], Eóin N. Molloy[1,3], Niklas Vockert[1], Jennifer Tremblay-Mercier[4], Jordana Remz[4], Alexa Pichet Binette[5,6], Sylvia Villeneuve[4,7], PREVENT-AD Research Group, Anne Maass[1,8]

[1] German Center for Neurodegenerative Diseases (DZNE), Magdeburg 39120, Germany
[2] Department of Neurobiology and Behavior, University of California, Irvine, USA
[3]Division of Nuclear Medicine, Department of Radiology & Nuclear Medicine, Faculty of Medicine, Otto von Guericke University Magdeburg, Magdeburg 39120, Germany
[4] Douglas Mental Health University Institute Research Centre, McGill University, Montréal H4H 1R3, Canada
[5] Clinical Memory Research, Faculty of Medicine, Lund University, Lund 223 62, Sweden
[6] Centre de Recherche de l'Institut Universitaire de Gériatrie de Montréal, H3W 1W5, Canada
[7] Department of Psychiatry, McGill University, Montréal H3A 1A1, Canada
[8] Institute for Biology, Otto von Guericke University Magdeburg, 39120 Magdeburg, Germany

**Please send correspondence to:**
Larissa Fischer,,
Anne Maass, Ph.D.

[a]A complete listing of the PREVENT-AD Research Group can be found at: https://preventad.loris.ca/acknowledgements/acknowledgements.php?DR=7.0&authors [b]Data used in preparation of this article were obtained from the Pre-symptomatic Evaluation of Experimental or Novel Treatments for Alzheimer's Disease (PREVENT-AD) program (https://www.centre-stopad.com/en/).

**Key Words:** Aging, Alzheimer's disease, fMRI, Functional Connectivity, Episodic Memory, *APOE*

#### Supplementary Figures:

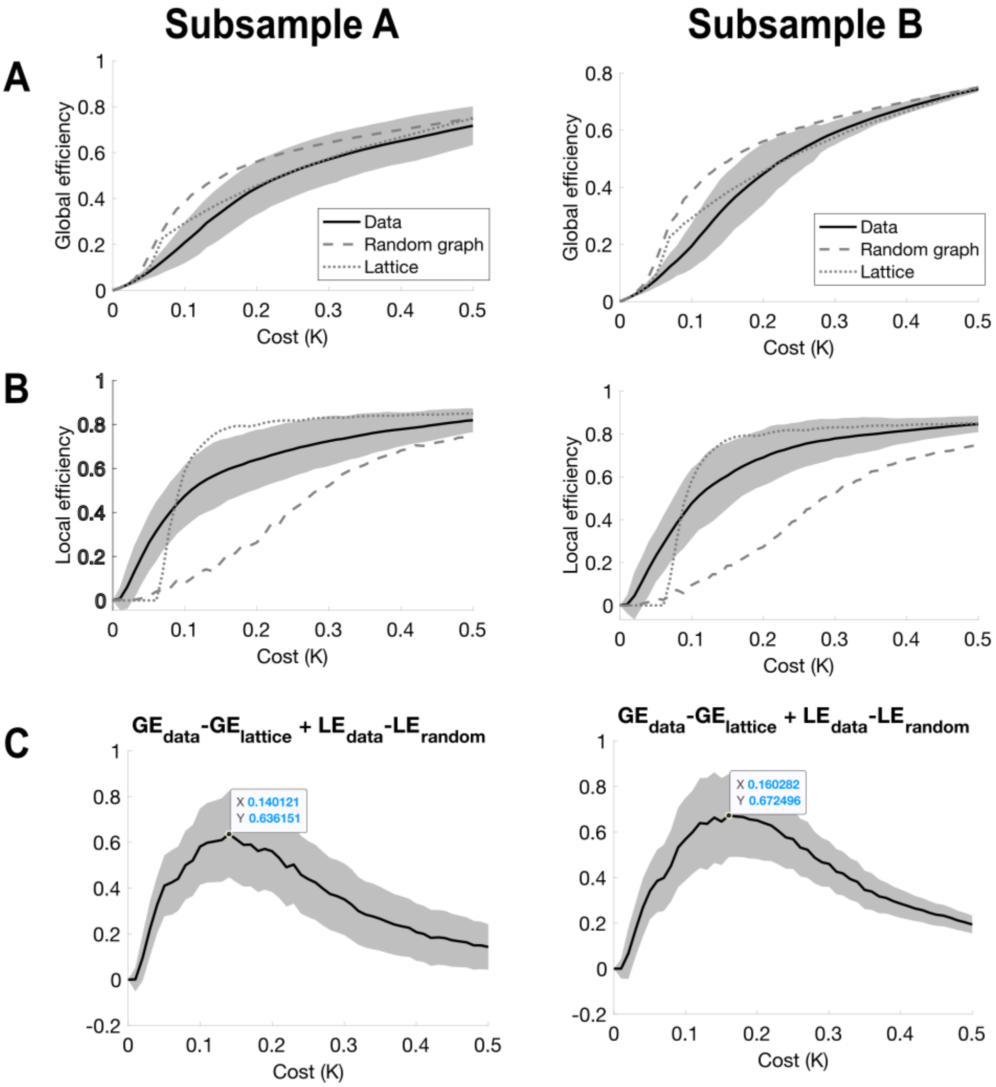


**Figure S1. Optimal cost threshold determination for subsample A and B. A)**. Global efficiency (GE) is plotted across varying cost thresholds, comparing how the real network (data) behaves relative to random and lattice structures. The random network acts as a baseline for randomization effects, while the lattice represents a highly ordered structure. **B)** Local efficiency (LE) is plotted across cost thresholds for the real, random, and lattice network. **C)** The optimal cost threshold is determined by maximising the difference between global and local efficiency values for real data compared to lattice and random networks, reflecting the network's small-world configuration. We used the formula *GE(Data) - GE(Lattice) + LE(Data) - LE(Random)*. The highest value on the y-axis is used as cost threshold, as it indicates the maximum divergence between the real and the modeled networks, with the real network's distinctive properties being most prominent. This resulted in an optimal cost threshold of 0.14 for subsample A and 0.16 for subsample B. See (Achard and Bullmore, 2007) for details.

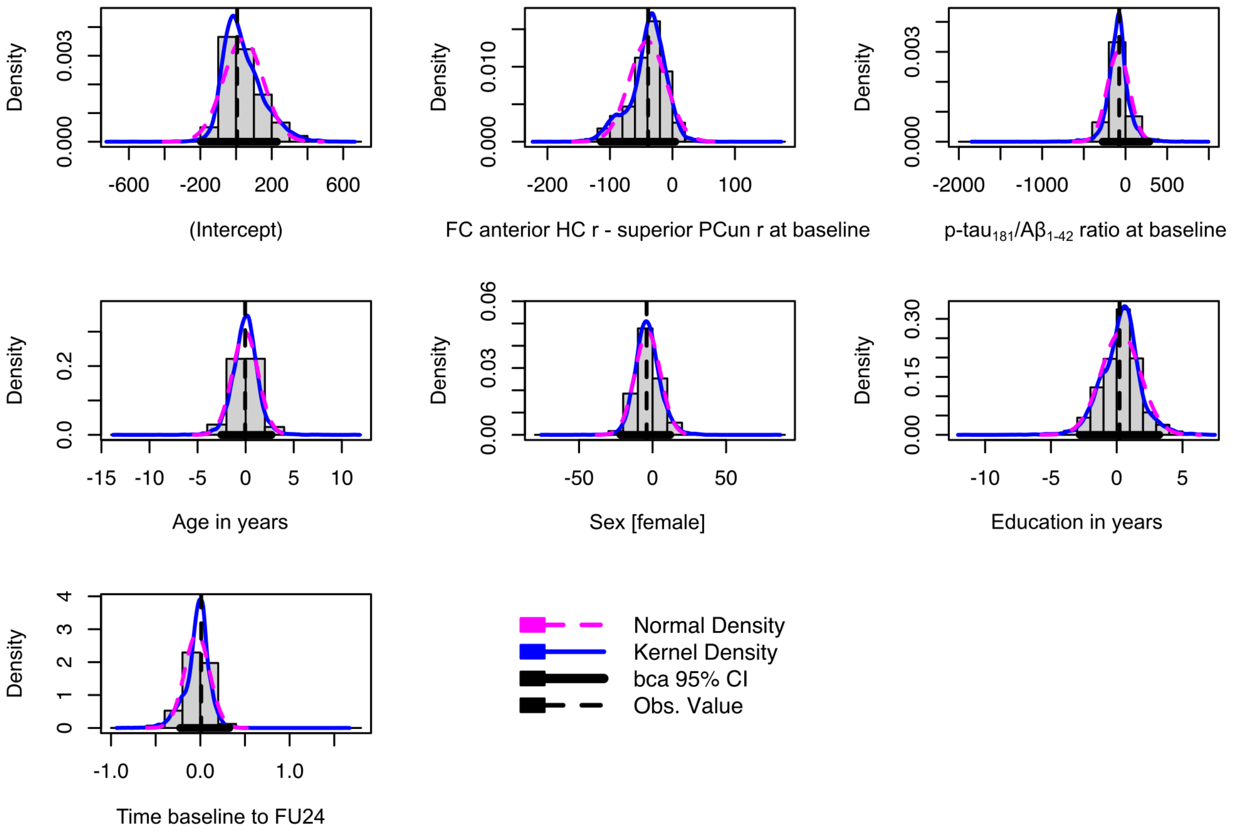


**Figure S2. Bootstrapped 95% confidence intervals for linear model of effect of connectivity on change in memory performance only in *APOE4* carriers**. For model, see Table S9. Change in RBANS episodic memory performance was used as dependent variable in *APOE4* carriers, functional connectivity at baseline, p-tau$_{181}$/A$\beta_{1\text{-}42}$ ratio at baseline, age at baseline, sex, education, and time between baseline and FU24 assessment were used as independent variables. RBANS = Repeatable Battery for Assessment of Neuropsychological Status. FC = functional connectivity. HC = hippocampus. r = right. PCun = precuneus. FU24 = follow-up 24 months after baseline. bca 95% CI = bias-corrected and accelerated 95% confidence interval. Obs. value = observed value.

#### Supplementary Tables:

##### Table S1. Medial temporal lobe -posteromedial network and visual Yeo network

| Network | ROI name | Brainnetome name (Left and Right Hemisphere) | Anatomical and modified Cyto-architectonic descriptions | Brainnetome coordinate L/ R MNI(X,Y,Z) |
|---|---|---|---|---|
| **MTL-PMC network** | | | | |
| | Anterior PRC | PhG_L(R)_6_1 | A35/36r, rostral area 35/36 | -27, -7, -34/ 28, -8, -33 |
| | Posterior PRC | PhG_L(R)_6_2 | A35/36c, caudal area 35/36 | -25, -25, -26/ 26, -23, -27 |
| | Lateral posterior PHC | PhG_L(R)_6_3 | TL, area TL (lateral PPHC, posterior parahippocampal gyrus) | -28, -32, -18/ 30, -30, -18 |
| | EC | PhG_L(R)_6_4 | A28/34, area 28/34 (EC, entorhinal cortex) | -19, -12, -30/ 19, -10, -30 |
| | Medial posterior PHC | PhG_L(R)_6_6 | TH, area TH (medial PPHC) | -17, -39, -10/ 19, -36, -11 |
| | Anterior hippocampus | Hipp_L(R)_2_1 | rHipp, rostral hippocampus | -22, -14, -19/ 22, -12, -20 |
| | Posterior hippocampus | Hipp_L(R)_2_2 | cHipp, caudal hippocampus | -28, -30, -10/ 29, -27, -10 |
| | PCC | CG_L(R)_7_1 | A23d, dorsal area 23 | -4, -39, 31/ 4, -37, 32 |
| | RSC | CG_L(R)_7_4 | A23v, ventral area 23 | -8, -47, 10/ 9, -44, 11 |
| | Lateral precuneus | PCun_L(R)_4_1 | A7m, medial area 7(PEp) | -5, -63, 51/ 6, -65, 51 |
| | Superior precuneus | PCun_L(R)_4_2 | A5m, medial area 5(PEm) | -8, -47, 57/ 7, -47, 58 |
| | Inferior precuneus | PCun_L(R)_4_3 | dmPOS, dorsomedial parietooccipital sulcus(PEr) | -12, -67, 25/ 16, -64, 25 |
| | Medial precuneus | PCun_L(R)_4_4 | A31, area 31 (Lc1) | -6, -55, 34/ 6, -54, 35 |
| | Superior mPFC | SFG_L(R)_7_7 | A10m, medial area 10 | -8, 56, 15/ 8, 58, 13 |
| | Inferior mPFC | OrG_L(R)_6_1 | A14m, medial area 14 | -7, 54, -7/ 6, 47, -7 |
| | Subgenual mPFC | CG_L(R)_7_7 | A32sg, subgenual area 32 | -4, 39, -2/ 5, 41, 6 |
| **Visual Yeo network** | | | | |
| | Fusiform gyrus | FuG_L(R)_3_2 | A37mv, medioventral area37 | -31, -64, -14/ 31, -62, -14 |
| | Posterior PRC | PhG_R_6_2 | A35/36c, caudal area 35/36 | 26, -23, -27 |
| | Lateral posterior PHC | PhG_L(R)_6_3 | TL, area TL (lateral PPHC, posterior parahippocampal gyrus) | -28, -32, -18/ 30, -30, -18 |
| | Medial posterior PHC | PhG_L(R)_6_6 | TH, area TH (medial PPHC) | -17, -39, -10/ 19, -36, -11 |

| | | | |
|---|---|---|---|
| Inferior parietal lobule | IPL_L(R)_6_1 | A39c, caudal area 39(PGp) | -34, -80, 29/ 45, -71, 20 |
| Inferior PCun | PCun_L(R)_4_3 | dmPOS, dorsomedial parietooccipital sulcus(PEr) | -12, -67, 25/ 16, -64, 25 |
| PCC | CG_R_7_4 | A23v, ventral area 23 | 9, -44, 11 |
| Medio-ventral occipital cortex | MVOcC_L(R)_5_1 | cLinG, caudal lingual gyrus | -11, -82, -11, 10, -85, -9 |
| Medio-ventral occipital cortex | MVOcC_L(R)_5_2 | rCunG, rostral cuneus gyrus | -5, -81, 10, 7, -76, 11 |
| Medio-ventral occipital cortex | MVOcC_L(R)_5_3 | cCunG, caudal cuneus gyrus | -6, -94, 1, 8, -90, 12 |
| Medio-ventral occipital cortex | MVOcC_L(R)_5_4 | rLinG, rostral lingual gyrus | -17, -60, -6, 18, -60, -7 |
| Medio-ventral occipital cortex | MVOcC_L(R)_5_5 | vmPOS,ventromedial parietooccipital sulcus | -13, -68, 12, 15, -63, 12 |
| Lateral occipital cortex | LOcC_L(R)_4_1 | mOccG, middle occipital gyrus | -31, -89, 11, 34, -86, 11 |
| Lateral occipital cortex | LOcC_R_4_2 | V5/MT+, area V5/MT+ | 48, -70, -1 |
| Lateral occipital cortex | LOcC_L(R)_4_3 | OPC, occipital polar cortex | -18, -99, 2, 22, -97, 4 |
| Lateral occipital cortex | LOcC_L(R)_4_4 | iOccG, inferior occipital gyrus | -30, -88, -12, 32, -85, -12 |
| Lateral occipital cortex | LOcC_L(R)_2_1 | msOccG, medial superior occipital gyrus | -11, -88, 31, 16, -85, 34 |
| Lateral occipital cortex | LOcC_L(R)_2_2 | lsOccG, lateral superior occipital gyrus | -22, -77, 36, 29, -75, 36 |

ROI = Region of interest. L = left. R = right. MTL = medial temporal lobe. PMC = posteromedial cortex. PRC = perirhinal cortex. PHC = parahippocampal cortex. EC = entorhinal cortex. PCC = Posterior Cingulate Cortex. RSC = Retrosplenial Cortex. mPFC = Medial Prefrontal Cortex.

#### Table S2. Subsample A: Linear model of effects of age on connectivity change

| | Change in FC lateral PHC r - posterior PRC l over one year | | | | | | | |
|---|---|---|---|---|---|---|---|---|
| *Predictors* | *Estimates* | *std. Error* | *std. Beta* | *standardized std. Error* | *CI* | *standardized CI* | *Statistic* | *p* |
| (Intercept) | 0.68 | 0.64 | 0.11 | 0.22 | -0.61 – 1.96 | -0.32 – 0.54 | 1.05 | 0.298 |
| Age in years | -0.02 | 0.01 | -0.34 | 0.11 | -0.04 – -0.01 | -0.55 – -0.13 | -3.15 | **0.002** |
| *APOE4* Group [carrier] | 0.01 | 0.07 | 0.03 | 0.22 | -0.12 – 0.14 | -0.42 – 0.48 | 0.13 | 0.895 |
| Sex [female] | -0.05 | 0.07 | -0.16 | 0.24 | -0.19 – 0.09 | -0.64 – 0.31 | -0.69 | 0.492 |
| Education in years | 0.01 | 0.01 | 0.07 | 0.10 | -0.01 – 0.02 | -0.13 – 0.27 | 0.70 | 0.485 |

| | | | | | | | | |
|---|---|---|---|---|---|---|---|---|
| Time baseline to FU12 | 0.00 | 0.00 | 0.14 | 0.10 | -0.00 – 0.00 | -0.06 – 0.35 | 1.42 | 0.159 |
| Observations | 93 | | | | | | | |
| $R^2$ / $R^2$ adjusted | 0.128 / 0.078 | | | | | | | |

Change in functional connectivity in amyloid and tau negative older adults was used as dependent variable, age at baseline, *APOE4* group, sex, education, and time between baseline and FU12 assessment were used as independent variables. FC = functional connectivity. PHC = Parahippocampal cortex. r = right. PRC = perirhinal cortex. l = left. CI = 95% confidence interval. FU12 = follow-up 12 months after baseline.

#### Table S3. Subsample A: Linear model of effects of age on connectivity change

| | Change in FC medial PHC r - PCC r over one year | | | | | | | |
|---|---|---|---|---|---|---|---|---|
| *Predictors* | *Estimates* | *std. Error* | *std. Beta* | *standardized std. Error* | *CI* | *standardized CI* | *Statistic* | *p* |
| (Intercept) | 0.86 | 0.46 | 0.07 | 0.22 | -0.06 – 1.78 | -0.36 – 0.49 | 1.86 | 0.066 |
| Age in years | -0.01 | 0.00 | -0.32 | 0.11 | -0.02 – -0.01 | -0.54 – -0.11 | -3.03 | **0.003** |
| *APOE4* Group [carrier] | -0.01 | 0.05 | -0.03 | 0.22 | -0.10 – 0.09 | -0.48 – 0.41 | -0.15 | 0.881 |
| Sex [female] | -0.02 | 0.05 | -0.08 | 0.24 | -0.12 – 0.08 | -0.55 – 0.39 | -0.33 | 0.745 |
| Education in years | 0.01 | 0.01 | 0.22 | 0.10 | 0.00 – 0.02 | 0.02 – 0.42 | 2.15 | **0.034** |
| Time baseline to FU12 | -0.00 | 0.00 | -0.04 | 0.10 | -0.00 – 0.00 | -0.24 – 0.16 | -0.43 | 0.670 |
| Observations | 93 | | | | | | | |
| $R^2$ / $R^2$ adjusted | 0.139 / 0.089 | | | | | | | |

Change in functional connectivity in amyloid and tau negative older adults was used as dependent variable, age at baseline, *APOE4* group, sex, education, and time between baseline and FU12 assessment were used as independent variables. FC = functional connectivity. PHC = Parahippocampal cortex. r = right. PCC = posterior cingulate cortex. CI = 95% confidence interval. FU12 = follow-up 12 months after baseline.

#### Table S4. Subsample A: Linear model of effects of age on connectivity change

| | Change in FC lateral PHC r - PCC l over one year | | | | | | | |
|---|---|---|---|---|---|---|---|---|
| *Predictors* | *Estimates* | *std. Error* | *std. Beta* | *standardized std. Error* | *CI* | *standardized CI* | *Statistic* | *p* |
| (Intercept) | 0.95 | 0.45 | 0.48 | 0.22 | 0.06 – 1.84 | 0.05 – 0.92 | 2.12 | **0.037** |

| | | | | | | | | |
|---|---|---|---|---|---|---|---|---|
| Age in years | -0.01 | 0.00 | -0.33 | 0.11 | -0.02 – -0.00 | -0.54 – -0.11 | -3.03 | **0.003** |
| *APOE4* Group [carrier] | -0.04 | 0.05 | -0.20 | 0.22 | -0.13 – 0.05 | -0.65 – 0.25 | -0.88 | 0.382 |
| Sex [female] | -0.12 | 0.05 | -0.58 | 0.24 | -0.22 – -0.02 | -1.06 – -0.11 | -2.43 | **0.017** |
| Education in years | 0.01 | 0.01 | 0.12 | 0.10 | -0.00 – 0.02 | -0.08 – 0.32 | 1.18 | 0.241 |
| Time baseline to FU12 | -0.00 | 0.00 | -0.02 | 0.10 | -0.00 – 0.00 | -0.22 – 0.18 | -0.20 | 0.839 |
| Observations | 93 | | | | | | | |
| $R^2$ / $R^2$ adjusted | 0.125 / 0.074 | | | | | | | |

Change in functional connectivity in amyloid and tau negative older adults was used as dependent variable, age at baseline, *APOE4* group, sex, education, and time between baseline and FU12 assessment were used as independent variables. FC = functional connectivity. PHC = Parahippocampal cortex. r = right. PCC = posterior cingulate cortex. l = left. CI = 95% confidence interval. FU12 = follow-up 12 months after baseline.

#### Table S5. Subsample B: Linear model of effects of AD pathology on connectivity change

| | Change in FC anterior HC r - superior PCun r over two years | | | | | | | |
|---|---|---|---|---|---|---|---|---|
| *Predictors* | *Estimates* | *std. Error* | *std. Beta* | *standardized std. Error* | *CI* | *standardized CI* | *Statistic* | *p* |
| (Intercept) | -0.07 | 0.43 | -0.22 | 0.22 | -0.93 – 0.80 | -0.66 – 0.22 | -0.15 | 0.879 |
| p-$tau_{181}$/$A\beta_{1-42}$ ratio at baseline | 4.31 | 1.03 | 0.51 | 0.12 | 2.25 – 6.37 | 0.27 – 0.76 | 4.19 | **<0.001** |
| Age in years | 0.00 | 0.00 | 0.10 | 0.12 | -0.00 – 0.01 | -0.13 – 0.34 | 0.89 | 0.375 |
| *APOE4* Group [carrier] | 0.01 | 0.05 | 0.03 | 0.25 | -0.10 – 0.11 | -0.48 – 0.54 | 0.11 | 0.909 |
| Sex [female] | 0.07 | 0.05 | 0.31 | 0.24 | -0.04 – 0.17 | -0.17 – 0.78 | 1.29 | 0.202 |
| Education in years | 0.00 | 0.01 | 0.04 | 0.12 | -0.01 – 0.02 | -0.19 – 0.27 | 0.34 | 0.734 |
| Time baseline to FU24 | -0.00 | 0.00 | -0.15 | 0.12 | -0.00 – 0.00 | -0.38 – 0.09 | -1.25 | 0.214 |
| Observations | 69 | | | | | | | |
| $R^2$ / $R^2$ adjusted | 0.288 / 0.220 | | | | | | | |

Change in functional connectivity was used as dependent variable, p-tau$_{181}$/A$\beta_{1-42}$ ratio at baseline, age at baseline, *APOE4* group, sex, education, and time between baseline and FU24 assessment were used as independent variables. AD = Alzheimer's disease. FC = functional connectivity. HC = hippocampus. r = right. PCun = precuneus. CI = 95% confidence interval. FU24 = follow-up 24 months after baseline.

#### Table S6. Subsample B: Linear model of effects of AD pathology on connectivity change

| | **Change in FC medial PHC r - superior PCun r over two years** | | | | | | | |
|---|---|---|---|---|---|---|---|---|
| *Predictors* | *Estimates* | *std. Error* | *std. Beta* | *standardized std. Error* | *CI* | *standardized CI* | *Statistic* | *p* |
| (Intercept) | -0.03 | 0.51 | -0.02 | 0.23 | -1.06 – 1.00 | -0.49 – 0.44 | -0.06 | 0.951 |
| p-tau$_{181}$/A$\beta_{1-42}$ ratio at baseline | 4.08 | 1.21 | 0.44 | 0.13 | 1.66 – 6.50 | 0.18 – 0.70 | 3.37 | **0.001** |
| Age in years | -0.00 | 0.01 | -0.01 | 0.12 | -0.01 – 0.01 | -0.26 – 0.23 | -0.11 | 0.912 |
| *APOE4* Group [carrier] | -0.01 | 0.06 | -0.05 | 0.27 | -0.14 – 0.12 | -0.59 – 0.49 | -0.20 | 0.843 |
| sex [female] | 0.01 | 0.06 | 0.06 | 0.25 | -0.10 – 0.13 | -0.44 – 0.56 | 0.24 | 0.809 |
| Education in years | -0.00 | 0.01 | -0.06 | 0.12 | -0.02 – 0.01 | -0.31 – 0.18 | -0.50 | 0.621 |
| Time baseline to FU24 | -0.00 | 0.00 | -0.02 | 0.12 | -0.00 – 0.00 | -0.27 – 0.22 | -0.18 | 0.855 |
| Observations | 69 | | | | | | | |
| $R^2$ / $R^2$ adjusted | 0.203 / 0.126 | | | | | | | |

Change in functional connectivity was used as dependent variable, p-tau$_{181}$/A$\beta_{1-42}$ ratio at baseline, age at baseline, *APOE4* group, sex, education, and time between baseline and FU24 assessment were used as independent variables. AD = Alzheimer's disease. FC = functional connectivity. PHC = parahippocampal cortex. r = right. PCun = precuneus. CI = 95% confidence interval. FU24 = follow-up 24 months after baseline.

#### Table S7. Subsample A: Linear model of effects of connectivity on change in memory

| Predictors | Change in RBANS episodic memory performance over two years | | | | | | | |
|---|---|---|---|---|---|---|---|---|
| | *Estimates* | *std. Error* | *std. Beta* | *standardized std. Error* | *CI* | *standardized CI* | *Statistic* | *p* |
| (Intercept) | 4.57 | 18.59 | -0.19 | 0.23 | -32.41 – 41.54 | -0.65 – 0.26 | 0.25 | 0.807 |
| FC RSC r - PCC r at baseline | 8.69 | 4.05 | 0.24 | 0.11 | 0.65 – 16.74 | 0.02 – 0.45 | 2.15 | **0.035** |
| Age in years | -0.12 | 0.19 | -0.07 | 0.11 | -0.49 – 0.26 | -0.30 – 0.16 | -0.62 | 0.540 |
| *APOE4* Group [carrier] | 2.24 | 1.88 | 0.29 | 0.24 | -1.49 – 5.98 | -0.19 – 0.76 | 1.19 | 0.236 |
| Sex [female] | 1.14 | 1.98 | 0.14 | 0.25 | -2.81 – 5.08 | -0.36 – 0.65 | 0.57 | 0.568 |
| Education in years | -0.23 | 0.23 | -0.11 | 0.11 | -0.69 – 0.23 | -0.32 – 0.11 | -0.99 | 0.324 |
| Time baseline to FU24 | 0.01 | 0.02 | 0.04 | 0.11 | -0.03 – 0.04 | -0.18 – 0.26 | 0.38 | 0.708 |
| Observations | 91 | | | | | | | |
| $R^2$ / $R^2$ adjusted | 0.079 / 0.014 | | | | | | | |

Change in RBANS episodic memory performance in amyloid and tau negative older adults was used as dependent variable, functional connectivity at baseline, age at baseline, *APOE4* group, sex, education, and time between baseline and FU24 assessment were used as independent variables. Due to multicollinearity, we left the interaction term of connectivity**APOE* out of the model. The interaction was not significant when included in the model. RBANS = Repeatable Battery for Assessment of Neuropsychological Status. FC = functional connectivity. RSC = retrosplenial cortex. r = right. PCC = posterior cingulate cortex. CI = 95% confidence interval. FU24 = follow-up 24 months after baseline.

#### Table S8. Subsample B: Linear model of effects of connectivity on change in memory

| Predictors | Change in RBANS episodic memory performance over two years | | | | | | | | |
|---|---|---|---|---|---|---|---|---|---|
| | *Estimates* | *std. Error* | *std. Beta* | *standardized std. Error* | *CI* | *standardized CI* | *Statistic* | *p* | *std. p* |
| (Intercept) | -16.74 | 23.18 | 0.00 | 0.26 | -63.12 – 29.64 | -0.52 – 0.52 | -0.72 | 0.473 | 0.997 |
| FC anterior HC r - superior PCun r at baseline | 9.41 | 10.59 | 0.14 | 0.16 | -11.77 – 30.60 | -0.18 – 0.47 | 0.89 | 0.378 | 0.378 |
| p-tau$_{181}$/A$\beta_{1-42}$ ratio at baseline | -36.03 | 62.35 | -0.09 | 0.15 | -160.79 – 88.73 | -0.38 – 0.21 | -0.58 | 0.566 | 0.566 |

| | | | | | | | | | |
|---|---|---|---|---|---|---|---|---|---|
| Age in years | 0.30 | 0.25 | 0.16 | 0.13 | -0.20 – 0.80 | -0.10 – 0.42 | 1.21 | 0.231 | 0.231 |
| *APOE4* Group [carrier] | 2.22 | 3.20 | 0.18 | 0.30 | -4.18 – 8.62 | -0.42 – 0.77 | 0.69 | 0.490 | 0.556 |
| Sex [female] | -0.39 | 2.94 | -0.04 | 0.27 | -6.28 – 5.50 | -0.59 – 0.51 | -0.13 | 0.895 | 0.895 |
| Education in years | 0.16 | 0.47 | 0.04 | 0.13 | -0.79 – 1.10 | -0.22 – 0.31 | 0.33 | 0.742 | 0.742 |
| Time baseline to FU24 | 0.00 | 0.03 | 0.01 | 0.13 | -0.05 – 0.06 | -0.25 – 0.28 | 0.11 | 0.912 | 0.912 |
| FC anterior HC r - superior PCun r at baseline × *APOE4* carrier [carrier] | -42.90 | 16.89 | -0.65 | 0.26 | -76.69 – -9.11 | -1.17 – -0.14 | -2.54 | **0.014** | **0.014** |
| Observations | 68 | | | | | | | | |
| $R^2$ / $R^2$ adjusted | 0.130 / 0.012 | | | | | | | | |

Change in RBANS episodic memory performance was used as dependent variable, functional connectivity at baseline, p-tau$_{181}$/A$\beta_{1\text{-}42}$ ratio at baseline, age at baseline, *APOE4* group, sex, education, time between baseline and FU24 assessment, and the interaction of connectivity**APOE* were used as independent variables. RBANS = Repeatable Battery for Assessment of Neuropsychological Status. FC = functional connectivity. HC = hippocampus. r = right. PCun = precuneus. CI = 95% confidence interval. FU24 = follow-up 24 months after baseline.

#### Table S9. Subsample B: Linear model of effects of connectivity on change in memory in *APOE4* carriers only

| | Change in RBANS episodic memory performance over two years | | | | | | | |
|---|---|---|---|---|---|---|---|---|
| *Predictors* | *Estimates* | *std. Error* | *std. Beta* | *standardized std. Error* | *CI* | *standardized CI* | *Statistic* | *p* |
| (Intercept) | 4.28 | 79.28 | 0.19 | 0.38 | -163.78 – 172.35 | -0.61 – 0.98 | 0.05 | 0.958 |
| FC anterior HC r - superior PCun r at baseline | -38.97 | 18.77 | -0.52 | 0.25 | -78.76 – 0.82 | -1.06 – 0.01 | -2.08 | **0.054** |
| p-tau$_{181}$/A$\beta_{1\text{-}42}$ ratio at baseline | -76.84 | 105.25 | -0.21 | 0.29 | -299.97 – 146.29 | -0.81 – 0.40 | -0.73 | 0.476 |
| Age in years | -0.01 | 1.10 | -0.00 | 0.24 | -2.35 – 2.32 | -0.52 – 0.51 | -0.01 | 0.990 |
| Sex [female] | -4.09 | 6.80 | -0.30 | 0.51 | -18.50 – 10.31 | -1.38 – 0.77 | -0.60 | 0.555 |
| Education in years | 0.21 | 1.05 | 0.06 | 0.27 | -2.00 – 2.43 | -0.52 – 0.63 | 0.21 | 0.840 |

| | | | | | | | | |
|---|---|---|---|---|---|---|---|---|
| Time baseline to FU24 | 0.01 | 0.09 | 0.02 | 0.24 | -0.17 – 0.19 | -0.49 – 0.54 | 0.10 | 0.920 |
| Observations | 23 | | | | | | | |
| $R^2$ / $R^2$ adjusted | 0.223 / -0.069 | | | | | | | |

Change in RBANS episodic memory performance was used as dependent variable in *APOE4* carriers, functional connectivity at baseline, p-tau$_{181}$/A$\beta_{1-42}$ ratio at baseline, age at baseline, sex, education, and time between baseline and FU24 assessment were used as independent variables. RBANS = Repeatable Battery for Assessment of Neuropsychological Status. FC = functional connectivity. HC = hippocampus. r = right. PCun = precuneus. CI = 95% confidence interval. FU24 = follow-up 24 months after baseline.

#### Table S10. Subsample A: Linear model of effects of change in connectivity on change in memory

| | Change in RBANS episodic memory performance over two years | | | | | | | |
|---|---|---|---|---|---|---|---|---|
| *Predictors* | *Estimates* | *std. Error* | *std. Beta* | *standardized std. Error* | *CI* | *standardized CI* | *Statistic* | *p* |
| (Intercept) | 6.07 | 18.41 | -0.09 | 0.22 | -30.54 – 42.68 | -0.52 – 0.35 | 0.33 | 0.743 |
| Change in FC RSC r - PCC r over two years | -1.00 | 3.33 | -0.03 | 0.11 | -7.62 – 5.62 | -0.25 – 0.19 | -0.30 | 0.764 |
| Age in years | -0.10 | 0.19 | -0.06 | 0.11 | -0.48 – 0.27 | -0.29 – 0.16 | -0.56 | 0.579 |
| Sex [female] | 0.91 | 2.03 | 0.12 | 0.26 | -3.13 – 4.95 | -0.40 – 0.63 | 0.45 | 0.655 |
| Education in years | -0.24 | 0.24 | -0.11 | 0.11 | -0.72 – 0.24 | -0.33 – 0.11 | -1.00 | 0.322 |
| Time baseline to FU24 | 0.01 | 0.02 | 0.05 | 0.11 | -0.03 – 0.05 | -0.17 – 0.27 | 0.48 | 0.633 |
| Observations | 91 | | | | | | | |
| $R^2$ / $R^2$ adjusted | 0.023 / -0.034 | | | | | | | |

Change in RBANS episodic memory performance in amyloid and tau negative older adults was used as dependent variable, change in functional connectivity, age at baseline, sex, education, and time between baseline and FU24 assessment were used as independent variables. RBANS = Repeatable Battery for Assessment of Neuropsychological Status. FC = functional connectivity. RSC = retrosplenial cortex. r = right. PCC = posterior cingulate cortex. CI = 95% confidence interval. FU24 = follow-up 24 months after baseline.

#### Table S11. Subsample A: Linear model of effects of change in connectivity and its interaction with *APOE* on change in memory

| | Change in RBANS episodic memory performance over two years | | | | | | | | |
|---|---|---|---|---|---|---|---|---|---|
| *Predictors* | *Estimates* | *std. Error* | *std. Beta* | *standardized std. Error* | *CI* | *standardized CI* | *Statistic* | *p* | *std. p* |
| (Intercept) | 1.00 | 19.13 | -0.17 | 0.24 | -37.05 – 39.06 | -0.65 – 0.30 | 0.05 | 0.958 | 0.464 |
| Change in FC RSC r - PCC r over two years | 0.54 | 3.83 | 0.02 | 0.13 | -7.09 – 8.17 | -0.24 – 0.27 | 0.14 | 0.888 | 0.888 |
| Age in years | -0.05 | 0.19 | -0.03 | 0.12 | -0.43 – 0.34 | -0.26 – 0.21 | -0.24 | 0.809 | 0.809 |
| *APOE4* Group [carrier] | 0.96 | 2.03 | 0.21 | 0.24 | -3.08 – 4.99 | -0.28 – 0.69 | 0.47 | 0.638 | 0.395 |
| Sex [female] | 1.23 | 2.06 | 0.16 | 0.26 | -2.86 – 5.33 | -0.36 – 0.68 | 0.60 | 0.551 | 0.551 |
| Education in years | -0.27 | 0.24 | -0.13 | 0.11 | -0.76 – 0.21 | -0.35 – 0.10 | -1.12 | 0.265 | 0.265 |
| Time baseline to FU24 | 0.01 | 0.02 | 0.06 | 0.11 | -0.03 – 0.05 | -0.16 – 0.29 | 0.57 | 0.567 | 0.567 |
| Change in FC RSC r - PCC r over two years × *APOE4* carrier [carrier] | -6.36 | 7.84 | -0.21 | 0.26 | -21.95 – 9.24 | -0.73 – 0.31 | -0.81 | 0.420 | 0.420 |
| Observations | 91 | | | | | | | | |
| $R^2$ / $R^2$ adjusted | 0.038 / -0.043 | | | | | | | | |

Change in RBANS episodic memory performance in amyloid and tau negative older adults was used as dependent variable, change in functional connectivity, age at baseline, *APOE4* group, sex, education, time between baseline and FU24 assessment, and the interaction of change in functional connectivity with *APOE4* group were used as independent variables. Due to multicollinearity, we left the interaction term of connectivity*age out of the model. The interaction was not significant when included in the model. RBANS = Repeatable Battery for Assessment of Neuropsychological Status. FC = functional connectivity. RSC = retrosplenial cortex. r = right. PCC = posterior cingulate cortex. CI = 95% confidence interval. FU24 = follow-up 24 months after baseline.

#### Table S12. Subsample B: Linear model of effects of change in connectivity on change in memory

| | Change in RBANS episodic memory performance over two years | | | | | | | |
|---|---|---|---|---|---|---|---|---|
| *Predictors* | *Estimates* | *std. Error* | *std. Beta* | *standardized std. Error* | *CI* | *standardized CI* | *Statistic* | *p* |
| (Intercept) | -16.81 | 22.60 | -0.02 | 0.23 | -61.98 – 28.37 | -0.48 – 0.45 | -0.74 | 0.460 |

| | | | | | | | | |
|---|---|---|---|---|---|---|---|---|
| Change in FC anterior HC r - superior PCun over two years | 2.93 | 6.44 | 0.06 | 0.13 | -9.95 – 15.81 | -0.20 – 0.31 | 0.46 | 0.650 |
| Age in years | 0.24 | 0.25 | 0.13 | 0.13 | -0.26 – 0.74 | -0.14 – 0.39 | 0.95 | 0.344 |
| Sex [female] | 0.26 | 2.97 | 0.02 | 0.28 | -5.69 – 6.20 | -0.53 – 0.58 | 0.09 | 0.931 |
| Education in years | 0.04 | 0.46 | 0.01 | 0.13 | -0.88 – 0.95 | -0.25 – 0.27 | 0.08 | 0.937 |
| Time baseline to FU24 | 0.01 | 0.03 | 0.04 | 0.13 | -0.05 – 0.06 | -0.22 – 0.31 | 0.33 | 0.745 |
| Observations | 68 | | | | | | | |
| $R^2$ / $R^2$ adjusted | 0.025 / -0.053 | | | | | | | |

Change in RBANS episodic memory performance was used as dependent variable, change in functional connectivity, age at baseline, sex, education, and time between baseline and FU24 assessment were used as independent variables. RBANS = Repeatable Battery for Assessment of Neuropsychological Status. FC = functional connectivity. HC = hippocampus. r = right. PCun = precuneus. CI = 95% confidence interval. FU24 = follow-up 24 months after baseline.

#### Table S13. Subsample B: Linear model of effects of change in connectivity and its interaction with *APOE* on change in memory

| | Change in RBANS episodic memory performance over two years | | | | | | | | |
|---|---|---|---|---|---|---|---|---|---|
| *Predictors* | *Estimates* | *std. Error* | *std. Beta* | *standardized std. Error* | *CI* | *standardized CI* | *Statistic* | *p* | *std. p* |
| (Intercept) | -14.68 | 24.66 | -0.05 | 0.27 | -64.01 – 34.66 | -0.60 – 0.49 | -0.60 | 0.554 | 0.853 |
| Change in FC anterior HC r - superior PCun over two years | -4.23 | 9.69 | -0.08 | 0.19 | -23.61 – 15.15 | -0.47 – 0.30 | -0.44 | 0.664 | 0.664 |
| p-$tau_{181}$/$A\beta_{1-42}$ ratio at baseline | -33.98 | 72.05 | -0.08 | 0.17 | -178.16 – 110.20 | -0.43 – 0.26 | -0.47 | 0.639 | 0.639 |
| Age in years | 0.28 | 0.26 | 0.15 | 0.14 | -0.25 – 0.80 | -0.13 – 0.42 | 1.06 | 0.292 | 0.292 |
| *APOE4* Group [carrier] | 1.23 | 3.31 | 0.06 | 0.30 | -5.39 – 7.84 | -0.55 – 0.67 | 0.37 | 0.712 | 0.844 |
| Sex [female] | 0.12 | 3.09 | 0.01 | 0.29 | -6.07 – 6.30 | -0.57 – 0.59 | 0.04 | 0.970 | 0.970 |
| Education in years | 0.04 | 0.49 | 0.01 | 0.14 | -0.94 – 1.02 | -0.26 – 0.28 | 0.08 | 0.938 | 0.938 |

| | | | | | | | | | |
|---|---|---|---|---|---|---|---|---|---|
| Time baseline to FU24 | 0.00 | 0.03 | 0.02 | 0.14 | -0.05 – 0.06 | -0.26 – 0.30 | 0.13 | 0.898 | 0.898 |
| Change in FC anterior HC r - superior PCun over two years × *APOE4* carrier [carrier] | 16.80 | 14.00 | 0.33 | 0.28 | -11.22 – 44.83 | -0.22 – 0.89 | 1.20 | 0.235 | 0.235 |
| Observations | 68 | | | | | | | | |
| $R^2$ / $R^2$ adjusted | 0.049 / -0.080 | | | | | | | | |

Change in RBANS episodic memory performance was used as dependent variable, change in functional connectivity, p-tau$_{181}$/Aβ$_{1-42}$ ratio at baseline, age at baseline, *APOE4* group, sex, education, time between baseline and FU24 assessment, and the interaction of change in functional connectivity with *APOE4* group were used as independent variables. Due to multicollinearity, we left the interaction term of connectivity* p-tau$_{181}$/Aβ$_{1-42}$ ratio at baseline out of the model. The interaction was not significant when included in the model. RBANS = Repeatable Battery for Assessment of Neuropsychological Status. FC = functional connectivity. HC = hippocampus. r = right. PCun = precuneus. CI = 95% confidence interval. FU24 = follow-up 24 months after baseline.

#### Table S14. Subsample A: Linear model of effects of connectivity on change in attention

| | **Change in RBANS attention performance over two years** | | | | | | | | |
|---|---|---|---|---|---|---|---|---|---|
| *Predictors* | *Estimates* | *std. Error* | *std. Beta* | *standardized std. Error* | *CI* | *standardized CI* | *Statistic* | *p* | *std. p* |
| (Intercept) | -39.90 | 36.64 | -0.10 | 0.23 | -112.80 – 33.01 | -0.56 – 0.36 | -1.09 | 0.279 | 0.674 |
| FC RSC r - PCC r at baseline | 10.63 | 8.52 | 0.17 | 0.13 | -6.31 – 27.58 | -0.10 – 0.43 | 1.25 | 0.215 | 0.215 |
| Age in years | 0.49 | 0.35 | 0.17 | 0.12 | -0.20 – 1.19 | -0.07 – 0.42 | 1.41 | 0.162 | 0.162 |
| *APOE4* Group [carrier] | 4.14 | 5.80 | 0.37 | 0.26 | -7.39 – 15.68 | -0.14 – 0.88 | 0.71 | 0.477 | 0.150 |
| Sex [female] | -0.08 | 3.51 | -0.01 | 0.26 | -7.07 – 6.91 | -0.52 – 0.51 | -0.02 | 0.982 | 0.982 |
| Education in years | -0.05 | 0.42 | -0.01 | 0.11 | -0.87 – 0.78 | -0.23 – 0.21 | -0.11 | 0.910 | 0.910 |
| Time baseline to FU24 | 0.01 | 0.04 | 0.02 | 0.11 | -0.07 – 0.08 | -0.20 – 0.23 | 0.14 | 0.890 | 0.890 |
| FC RSC r - PCC r at baseline × *APOE4* carrier [carrier] | 2.79 | 16.80 | 0.04 | 0.26 | -30.64 – 36.23 | -0.48 – 0.57 | 0.17 | 0.868 | 0.868 |

| | |
|---|---|
| Observations | 88 |
| $R^2$ / $R^2$ adjusted | 0.079 / -0.001 |

Change in RBANS attention performance in amyloid and tau negative older adults was used as dependent variable, functional connectivity at baseline, age at baseline, *APOE4* group, sex, education, time between baseline and FU24 assessment and the interaction of FC**APOE4* group were used as independent variables. RBANS = Repeatable Battery for Assessment of Neuropsychological Status. FC = functional connectivity. RSC = retrosplenial cortex. r = right. PCC = posterior cingulate cortex. CI = 95% confidence interval. FU24 = follow-up 24 months after baseline.

#### Table S15. Subsample B: Linear model of effects of connectivity on change in attention

| | **Change in RBANS attention performance over two years** | | | | | | | | |
|---|---|---|---|---|---|---|---|---|---|
| *Predictors* | *Estimates* | *std. Error* | *std. Beta* | *standardized std. Error* | *CI* | *standardized CI* | *Statistic* | *p* | *std. p* |
| (Intercept) | -49.45 | 26.38 | -0.18 | 0.24 | -102.23 – 3.34 | -0.65 – 0.30 | -1.87 | **0.066** | 0.457 |
| FC anterior HC r - superior PCun r at baseline | 5.49 | 12.05 | 0.07 | 0.15 | -18.62 – 29.61 | -0.23 – 0.36 | 0.46 | 0.650 | 0.650 |
| p-$tau_{181}$/$A\beta_{1-42}$ ratio at baseline | -142.98 | 70.95 | -0.27 | 0.14 | -284.96 – -1.00 | -0.55 – -0.00 | -2.02 | **0.048** | **0.048** |
| Age in years | 0.97 | 0.28 | 0.41 | 0.12 | 0.40 – 1.54 | 0.17 – 0.65 | 3.39 | **0.001** | **0.001** |
| *APOE4* Group [carrier] | 8.30 | 3.64 | 0.61 | 0.27 | 1.01 – 15.59 | 0.06 – 1.15 | 2.28 | **0.026** | **0.029** |
| Sex [female] | -0.23 | 3.35 | -0.02 | 0.25 | -6.93 – 6.47 | -0.52 – 0.49 | -0.07 | 0.945 | 0.945 |
| Education in years | 1.09 | 0.54 | 0.24 | 0.12 | 0.01 – 2.16 | 0.00 – 0.49 | 2.02 | **0.048** | **0.048** |
| Time baseline to FU24 | -0.03 | 0.03 | -0.13 | 0.12 | -0.09 – 0.03 | -0.37 – 0.12 | -1.04 | 0.302 | 0.302 |
| FC anterior HC r - superior PCun r at baseline × *APOE4* carrier [carrier] | -24.85 | 19.22 | -0.30 | 0.24 | -63.30 – 13.60 | -0.78 – 0.17 | -1.29 | 0.201 | 0.201 |
| Observations | 68 | | | | | | | | |
| $R^2$ / $R^2$ adjusted | 0.271 / 0.172 | | | | | | | | |

Change in RBANS attention performance was used as dependent variable, functional connectivity at baseline, age at baseline, *APOE4* group, sex, education, time between baseline and FU24 assessment and the interaction of

FC**APOE4* group were used as independent variables. RBANS = Repeatable Battery for Assessment of Neuropsychological Status. FC = functional connectivity. HC = hippocampus. r = right. PCun = precuneus. CI = 95% confidence interval. FU24 = follow-up 24 months after baseline.

#### Table S16. Subsample A: Linear model of effects of change in connectivity on change in attention

| | Change in RBANS attention performance over two years | | | | | | | |
|---|---|---|---|---|---|---|---|---|
| *Predictors* | *Estimates* | *std. Error* | *std. Beta* | *standardized std. Error* | *CI* | *standardized CI* | *Statistic* | *p* |
| (Intercept) | -33.86 | 35.82 | 0.05 | 0.22 | -105.11 – 37.40 | -0.38 – 0.48 | -0.95 | 0.347 |
| Change in FC RSC r - PCC r over two years | 5.97 | 5.77 | 0.12 | 0.11 | -5.51 – 17.46 | -0.11 – 0.34 | 1.04 | 0.304 |
| Age in years | 0.47 | 0.32 | 0.17 | 0.11 | -0.17 – 1.11 | -0.06 – 0.39 | 1.45 | 0.151 |
| Sex [female] | -1.00 | 3.51 | -0.07 | 0.26 | -7.97 – 5.98 | -0.58 – 0.44 | -0.28 | 0.777 |
| Education in years | -0.12 | 0.42 | -0.03 | 0.11 | -0.95 – 0.72 | -0.25 – 0.19 | -0.28 | 0.781 |
| Time baseline to FU24 | 0.01 | 0.04 | 0.03 | 0.11 | -0.07 – 0.09 | -0.19 – 0.24 | 0.24 | 0.808 |
| Observations | 88 | | | | | | | |
| $R^2$ / $R^2$ adjusted | 0.041 / -0.018 | | | | | | | |

Change in RBANS attention performance in amyloid and tau negative older adults was used as dependent variable, change in functional connectivity, age at baseline, sex, education, and time between baseline and FU24 assessment were used as independent variables. RBANS = Repeatable Battery for Assessment of Neuropsychological Status. FC = functional connectivity. RSC = retrosplenial cortex. r = right. PCC = posterior cingulate cortex. CI = 95% confidence interval. FU24 = follow-up 24 months after baseline.

#### Table S17. Subsample A: Linear model of effects of change in connectivity and its interaction with *APOE* on change in attention

| | Change in RBANS attention performance over two years | | | | | | | | |
|---|---|---|---|---|---|---|---|---|---|
| *Predictors* | *Estimates* | *std. Error* | *std. Beta* | *standardized std. Error* | *CI* | *standardized CI* | *Statistic* | *p* | *std. p* |

| | | | | | | | | | |
|---|---|---|---|---|---|---|---|---|---|
| (Intercept) | -42.08 | 36.61 | -0.05 | 0.23 | -114.94 – 30.78 | -0.52 – 0.41 | -1.15 | 0.254 | 0.819 |
| Change in FC RSC r - PCC r over two years | 6.53 | 6.62 | 0.13 | 0.13 | -6.64 – 19.71 | -0.13 – 0.38 | 0.99 | 0.327 | 0.327 |
| Age in years | 0.55 | 0.34 | 0.20 | 0.12 | -0.12 – 1.22 | -0.04 – 0.43 | 1.65 | 0.104 | 0.104 |
| *APOE4* Group [carrier] | 3.88 | 3.59 | 0.30 | 0.25 | -3.27 – 11.03 | -0.19 – 0.79 | 1.08 | 0.283 | 0.229 |
| Sex [female] | -0.58 | 3.55 | -0.04 | 0.26 | -7.65 – 6.49 | -0.56 – 0.47 | -0.16 | 0.870 | 0.870 |
| Education in years | -0.20 | 0.43 | -0.05 | 0.11 | -1.05 – 0.65 | -0.28 – 0.17 | -0.47 | 0.641 | 0.641 |
| Time baseline to FU24 | 0.01 | 0.04 | 0.04 | 0.11 | -0.06 – 0.09 | -0.18 – 0.26 | 0.34 | 0.734 | 0.734 |
| Change in FC RSC r - PCC r over two years × *APOE4* Group [carrier] | -2.06 | 13.71 | -0.04 | 0.26 | -29.35 – 25.23 | -0.57 – 0.49 | -0.15 | 0.881 | 0.881 |
| Observations | 88 | | | | | | | | |
| $R^2$ / $R^2$ adjusted | 0.058 / -0.024 | | | | | | | | |

Change in RBANS attention performance in amyloid and tau negative older adults was used as dependent variable, change in functional connectivity, age at baseline, *APOE4* group, sex, education, time between baseline and FU24 assessment, and the interaction of change in functional connectivity with *APOE4* group were used as independent variables. Due to multicollinearity, we left the interaction term of connectivity*age out of the model. The interaction was not significant when included in the model. RBANS = Repeatable Battery for Assessment of Neuropsychological Status. FC = functional connectivity. RSC = retrosplenial cortex. r = right. PCC = posterior cingulate cortex. CI = 95% confidence interval. FU24 = follow-up 24 months after baseline.

#### Table S18. Subsample B: Linear model of effects of change in connectivity on change in attention

| | **Change in RBANS attention performance over two years** | | | | | | | |
|---|---|---|---|---|---|---|---|---|
| *Predictors* | *Estimates* | *std. Error* | *std. Beta* | *standardized std. Error* | *CI* | *standardized CI* | *Statistic* | *p* |
| (Intercept) | -39.95 | 25.53 | -0.01 | 0.21 | -90.98 – 11.09 | -0.43 – 0.41 | -1.56 | 0.123 |

| | | | | | | | | |
|---|---|---|---|---|---|---|---|---|
| Change in FC anterior HC r - superior PCun r over two years | -8.99 | 7.28 | -0.14 | 0.12 | -23.54 – 5.56 | -0.38 – 0.09 | -1.24 | 0.221 |
| Age in years | 0.80 | 0.28 | 0.34 | 0.12 | 0.23 – 1.36 | 0.10 – 0.58 | 2.83 | **0.006** |
| Sex [female] | 0.15 | 3.36 | 0.01 | 0.25 | -6.57 – 6.87 | -0.49 – 0.52 | 0.04 | **0.964** |
| Education in years | 1.22 | 0.52 | 0.27 | 0.12 | 0.18 – 2.26 | 0.04 – 0.51 | 2.35 | **0.022** |
| Time baseline to FU24 | -0.04 | 0.03 | -0.15 | 0.12 | -0.10 – 0.02 | -0.39 – 0.09 | -1.26 | 0.211 |
| Observations | 68 | | | | | | | |
| $R^2$ / $R^2$ adjusted | 0.195 / 0.131 | | | | | | | |

Change in RBANS attention performance was used as dependent variable, change in functional connectivity, age at baseline, sex, education, and time between baseline and FU24 assessment were used as independent variables. RBANS = Repeatable Battery for Assessment of Neuropsychological Status. FC = functional connectivity. HC = hippocampus. r = right. PCun = precuneus. CI = 95% confidence interval. FU24 = follow-up 24 months after baseline.

#### Table S19. Subsample B: Linear model of effects of change in connectivity and its interaction with *APOE* on change in attention

| | Change in RBANS attention performance over two years | | | | | | | | |
|---|---|---|---|---|---|---|---|---|---|
| *Predictors* | *Estimates* | *std. Error* | *std. Beta* | *standardized std. Error* | *CI* | *standardized CI* | *Statistic* | *p* | *std. p* |
| (Intercept) | -50.13 | 26.60 | -0.25 | 0.24 | -103.36 – 3.09 | -0.73 – 0.22 | -1.88 | **0.064** | 0.289 |
| Change in FC anterior HC r - superior PCun r over two years | -17.60 | 10.45 | -0.28 | 0.17 | -38.51 – 3.31 | -0.61 – 0.05 | -1.68 | 0.097 | 0.097 |
| $p\text{-}tau_{181}/A\beta_{1-42}$ ratio at baseline | -125.86 | 77.73 | -0.24 | 0.15 | -281.40 – 29.68 | -0.54 – 0.06 | -1.62 | 0.111 | 0.111 |
| Age in years | 1.00 | 0.28 | 0.42 | 0.12 | 0.43 – 1.57 | 0.18 – 0.66 | 3.52 | **0.001** | **0.001** |
| *APOE4* Group [carrier] | 8.47 | 3.57 | 0.57 | 0.26 | 1.33 – 15.61 | 0.05 – 1.10 | 2.37 | **0.021** | **0.033** |
| Sex [female] | 0.61 | 3.34 | 0.05 | 0.25 | -6.06 – 7.29 | -0.46 – 0.55 | 0.18 | 0.855 | 0.855 |

| | | | | | | | | | |
|---|---|---|---|---|---|---|---|---|---|
| Education in years | 1.04 | 0.53 | 0.23 | 0.12 | -0.01 – 2.10 | -0.00 – 0.47 | 1.97 | 0.053 | 0.053 |
| Time baseline to FU24 | -0.04 | 0.03 | -0.14 | 0.12 | -0.10 – 0.03 | -0.38 – 0.10 | -1.17 | 0.245 | 0.245 |
| Change in FC anterior HC r - superior PCun r over two years × *APOE4* Group [carrier] | 24.01 | 15.11 | 0.38 | 0.24 | -6.22 – 54.25 | -0.10 – 0.86 | 1.59 | 0.117 | 0.117 |
| Observations | 68 | | | | | | | | |
| $R^2$ / $R^2$ adjusted | 0.284 / 0.187 | | | | | | | | |

Change in RBANS attention performance was used as dependent variable, change in functional connectivity, p-$tau_{181}$/$A\beta_{1-42}$ ratio at baseline, age at baseline, *APOE4* group, sex, education, time between baseline and FU24 assessment, and the interaction of change in functional connectivity with *APOE4* group were used as independent variables. Due to multicollinearity, we left the interaction term of connectivity* p-$tau_{181}$/$A\beta_{1-42}$ ratio at baseline out of the model. The interaction was not significant when included in the model. RBANS = Repeatable Battery for Assessment of Neuropsychological Status. FC = functional connectivity. HC = hippocampus. r = right. PCun = precuneus. CI = 95% confidence interval. FU24 = follow-up 24 months after baseline.

#### Table S20. Post-hoc comparisons of change in hippocampus volume over time

| **Subsample** | **Baseline to FU12** | **Baseline to FU24** | **FU12 to FU24** |
|---|---|---|---|
| A | 0.007 | <0.001 | <0.001 |
| B | 0.117 | <0.001 | 0.009 |

Displayed are bonferroni-corrected p-values. FU12 = follow-up 12 months after baseline. FU24 = follow-up 24 months after baseline.

#### Table S21. Subsample B: Linear model of effects of AD pathology on change in hippocampus volume

| | **Change in hippocampus volume over two years** | | | | | | | |
|---|---|---|---|---|---|---|---|---|
| *Predictors* | *Estimates* | *std. Error* | *std. Beta* | *standardized std. Error* | *CI* | *standardized CI* | *Statistic* | *p* |
| (Intercept) | 13.41 | 4.16 | 0.01 | 0.23 | 5.09 – 21.73 | -0.44 – 0.47 | 3.22 | **0.002** |
| p-$tau_{181}$/$A\beta_{1-42}$ ratio at baseline | -12.54 | 9.89 | -0.16 | 0.13 | -32.30 – 7.22 | -0.41 – 0.09 | -1.27 | 0.209 |
| Age in years | -0.07 | 0.04 | -0.20 | 0.12 | -0.16 – 0.01 | -0.44 – 0.04 | -1.69 | 0.097 |

| | | | | | | | | |
|---|---|---|---|---|---|---|---|---|
| *APOE4* Group [carrier] | -0.27 | 0.52 | -0.14 | 0.26 | -1.31 – 0.77 | -0.66 – 0.39 | -0.52 | 0.603 |
| Sex [female] | 0.09 | 0.49 | 0.05 | 0.24 | -0.88 – 1.06 | -0.44 – 0.53 | 0.19 | 0.850 |
| Education in years | 0.09 | 0.08 | 0.13 | 0.12 | -0.07 – 0.25 | -0.10 – 0.37 | 1.11 | 0.269 |
| Time baseline to FU24 | -0.01 | 0.01 | -0.34 | 0.12 | -0.02 – -0.00 | -0.57 – -0.10 | -2.82 | **0.007** |
| Observations | 69 | | | | | | | |
| $R^2$ / $R^2$ adjusted | 0.250 / 0.177 | | | | | | | |

Change in hippocampal volume was used as dependent variable, p-tau181/Aβ1-42 ratio at baseline, age at baseline, *APOE4* group, sex, education, and time between baseline and FU24 assessment were used as independent variables. AD = Alzheimer's disease. CI = 95% confidence interval. FU24 = follow-up 24 months after baseline.

#### Table S22. Subsample A: Linear model of effects of age and change in hippocampal volume on connectivity change

| | **Change in FC lateral PHC r - posterior PRC l over one year** | | | | | | | |
|---|---|---|---|---|---|---|---|---|
| *Predictors* | *Estimates* | *std. Error* | *std. Beta* | *standardized std. Error* | *CI* | *standardized CI* | *Statistic* | *p* |
| (Intercept) | 0.68 | 0.64 | 0.14 | 0.21 | -0.59 – 1.95 | -0.29 – 0.57 | 1.07 | 0.288 |
| Age in years | -0.02 | 0.01 | -0.36 | 0.11 | -0.04 – -0.01 | -0.57 – -0.15 | -3.37 | **0.001** |
| *APOE4* Group [carrier] | -0.01 | 0.07 | -0.02 | 0.22 | -0.14 – 0.12 | -0.47 – 0.42 | -0.11 | 0.912 |
| sex [female] | -0.05 | 0.07 | -0.18 | 0.24 | -0.19 – 0.09 | -0.65 – 0.29 | -0.76 | 0.448 |
| Education in years | 0.01 | 0.01 | 0.12 | 0.10 | -0.01 – 0.03 | -0.09 – 0.32 | 1.13 | 0.260 |
| Time baseline to FU12 | 0.00 | 0.00 | 0.15 | 0.10 | -0.00 – 0.00 | -0.05 – 0.35 | 1.46 | 0.148 |
| HC volume change over one year | -0.04 | 0.02 | -0.18 | 0.10 | -0.08 – 0.01 | -0.39 – 0.02 | -1.75 | 0.084 |
| Observations | 93 | | | | | | | |
| $R^2$ / $R^2$ adjusted | 0.158 / 0.100 | | | | | | | |

Change in functional connectivity in amyloid and tau negative older adults was used as dependent variable, age at baseline, *APOE4* group, sex, education, time between baseline and FU12 assessment, and change in hippocampus volume from baseline to FU12 were used as independent variables. FC = functional connectivity. PHC = Parahippocampal cortex. r = right. PRC = perirhinal cortex. l = left. CI = 95% confidence interval. FU12 = follow-up 12 months after baseline. HC = hippocampus.

#### Table S23. Subsample A: Linear model of effects of age and change in hippocampal volume on connectivity change

| | Change in FC medial PHC r - PCC r over one year | | | | | | | |
|---|---|---|---|---|---|---|---|---|
| *Predictors* | *Estimates* | *std. Error* | *std. Beta* | *standardized std. Error* | *CI* | *standardized CI* | *Statistic* | *p* |
| (Intercept) | 0.86 | 0.46 | 0.08 | 0.22 | -0.06 – 1.78 | -0.35 – 0.51 | 1.87 | 0.066 |
| Age in years | -0.02 | 0.00 | -0.34 | 0.11 | -0.03 – -0.01 | -0.55 – -0.12 | -3.12 | **0.002** |
| *APOE4* Group [carrier] | -0.01 | 0.05 | -0.06 | 0.23 | -0.11 – 0.08 | -0.51 – 0.38 | -0.28 | 0.779 |
| sex [female] | -0.02 | 0.05 | -0.09 | 0.24 | -0.12 – 0.08 | -0.56 – 0.39 | -0.36 | 0.718 |
| Education in years | 0.01 | 0.01 | 0.24 | 0.10 | 0.00 – 0.03 | 0.03 – 0.45 | 2.32 | **0.023** |
| Time baseline to FU12 | -0.00 | 0.00 | -0.04 | 0.10 | -0.00 – 0.00 | -0.24 – 0.16 | -0.41 | 0.680 |
| HC volume change over one year | -0.02 | 0.02 | -0.10 | 0.10 | -0.05 – 0.02 | -0.31 – 0.11 | -0.95 | 0.344 |
| Observations | 93 | | | | | | | |
| $R^2$ / $R^2$ adjusted | 0.148 / 0.088 | | | | | | | |

Change in functional connectivity in amyloid and tau negative older adults was used as dependent variable, age at baseline, *APOE4* group, sex, education, time between baseline and FU12 assessment, and change in hippocampus volume from baseline to FU12 were used as independent variables. FC = functional connectivity. PHC = Parahippocampal cortex. r = right. PCC = posterior cingulate cortex. CI = 95% confidence interval. FU12 = follow-up 12 months after baseline. HC = hippocampus.

#### Table S24. Subsample A: Linear model of effects of age and change in hippocampal volume on connectivity change

| | Change in FC lateral PHC r - PCC l over one year | | | | | | | |
|---|---|---|---|---|---|---|---|---|
| *Predictors* | *Estimates* | *std. Error* | *std. Beta* | *standardized std. Error* | *CI* | *standardized CI* | *Statistic* | *p* |
| (Intercept) | 0.95 | 0.45 | 0.49 | 0.22 | 0.06 – 1.84 | 0.06 – 0.93 | 2.12 | **0.037** |
| Age in years | -0.01 | 0.00 | -0.33 | 0.11 | -0.02 – -0.01 | -0.55 – -0.12 | -3.06 | **0.003** |
| *APOE4* Group [carrier] | -0.04 | 0.05 | -0.22 | 0.23 | -0.14 – 0.05 | -0.67 – 0.24 | -0.95 | 0.345 |
| sex [female] | -0.12 | 0.05 | -0.59 | 0.24 | -0.22 – -0.02 | -1.06 – -0.11 | -2.45 | **0.017** |
| Education in years | 0.01 | 0.01 | 0.14 | 0.11 | -0.00 – 0.02 | -0.07 – 0.35 | 1.29 | 0.201 |
| Time baseline to FU12 | -0.00 | 0.00 | -0.02 | 0.10 | -0.00 – 0.00 | -0.22 – 0.18 | -0.19 | 0.846 |
| HC volume change over one year | -0.01 | 0.02 | -0.06 | 0.11 | -0.04 – 0.02 | -0.27 – 0.15 | -0.58 | 0.561 |
| Observations | 93 | | | | | | | |
| $R^2$ / $R^2$ adjusted | 0.128 / 0.067 | | | | | | | |

Change in functional connectivity in amyloid and tau negative older adults was used as dependent variable, age at baseline, *APOE4* group, sex, education, time between baseline and FU12 assessment, and change in hippocampus volume from baseline to FU12 were used as independent variables. FC = functional connectivity. PHC = Parahippocampal cortex. r = right. PCC = posterior cingulate cortex. l = left. CI = 95% confidence interval. FU12 = follow-up 12 months after baseline. HC = hippocampus.

#### Table S25. Subsample B: Linear model of effects of AD pathology and change in hippocampal volume on connectivity change

| | Change in FC anterior HC r - superior PCun r over two years | | | | | | | |
|---|---|---|---|---|---|---|---|---|
| *Predictors* | *Estimates* | *std. Error* | *std. Beta* | *standardized std. Error* | *CI* | *standardized CI* | *Statistic* | *p* |
| (Intercept) | 0.08 | 0.47 | -0.22 | 0.22 | -0.86 – 1.02 | -0.66 – 0.22 | 0.18 | 0.861 |

| | | | | | | | | |
|---|---|---|---|---|---|---|---|---|
| p-tau181/Aβ1-42 ratio at baseline | 4.17 | 1.04 | 0.50 | 0.12 | 2.08 – 6.26 | 0.25 – 0.75 | 3.99 | **<0.001** |
| Age in years | 0.00 | 0.00 | 0.08 | 0.12 | -0.01 – 0.01 | -0.16 – 0.32 | 0.70 | 0.488 |
| *APOE4* Group [carrier] | 0.00 | 0.05 | 0.02 | 0.26 | -0.11 – 0.11 | -0.50 – 0.53 | 0.06 | 0.953 |
| sex [female] | 0.07 | 0.05 | 0.31 | 0.24 | -0.04 – 0.17 | -0.17 – 0.79 | 1.30 | 0.197 |
| Education in years | 0.00 | 0.01 | 0.05 | 0.12 | -0.01 – 0.02 | -0.18 – 0.29 | 0.45 | 0.651 |
| Time baseline to FU12 | -0.00 | 0.00 | -0.18 | 0.12 | -0.00 – 0.00 | -0.43 – 0.07 | -1.46 | 0.149 |
| HC volume change over two years | -0.01 | 0.01 | -0.10 | 0.12 | -0.04 – 0.02 | -0.35 – 0.14 | -0.84 | 0.407 |
| Observations | 69 | | | | | | | |
| $R^2$ / $R^2$ adjusted | 0.297 / 0.216 | | | | | | | |

Change in functional connectivity was used as dependent variable, p-tau181/Aβ1-42 ratio at baseline, age at baseline, *APOE4* group, sex, education, time between baseline and FU12 assessment, and change in hippocampus volume from baseline to FU12 were used as independent variables. AD = Alzheimer's disease. FC = functional connectivity. HC = hippocampus. r = right. PCun = precuneus. CI = 95% confidence interval. FU24 = follow-up 24 months after baseline.

#### Table S26. Subsample B: Linear model of effects of AD pathology and change in hippocampal volume on connectivity change

| | Change in FC medial PHC r - superior PCun r over two years | | | | | | | |
|---|---|---|---|---|---|---|---|---|
| *Predictors* | *Estimates* | *std. Error* | *std. Beta* | *standardized std. Error* | *CI* | *standardized CI* | *Statistic* | *p* |
| (Intercept) | -0.08 | 0.56 | -0.03 | 0.24 | -1.19 – 1.03 | -0.50 – 0.45 | -0.14 | 0.889 |
| p-tau181/Aβ1-42 ratio at baseline | 4.12 | 1.24 | 0.44 | 0.13 | 1.65 – 6.59 | 0.18 – 0.71 | 3.33 | **0.001** |
| Age in years | -0.00 | 0.01 | -0.01 | 0.13 | -0.01 – 0.01 | -0.26 – 0.25 | -0.05 | 0.958 |
| *APOE4* Group [carrier] | -0.01 | 0.06 | -0.05 | 0.27 | -0.14 – 0.12 | -0.59 – 0.50 | -0.18 | 0.858 |

| | | | | | | | | |
|---|---|---|---|---|---|---|---|---|
| sex [female] | 0.01 | 0.06 | 0.06 | 0.25 | -0.11 – 0.13 | -0.45 – 0.57 | 0.24 | 0.814 |
| Education in years | -0.01 | 0.01 | -0.06 | 0.12 | -0.02 – 0.01 | -0.31 – 0.18 | -0.52 | 0.607 |
| Time baseline to FU24 | -0.00 | 0.00 | -0.01 | 0.13 | -0.00 – 0.00 | -0.28 – 0.25 | -0.10 | 0.917 |
| HC volume change over two years | 0.00 | 0.02 | 0.03 | 0.13 | -0.03 – 0.03 | -0.24 – 0.29 | 0.20 | 0.842 |
| Observations | 69 | | | | | | | |
| $R^2$ / $R^2$ adjusted | 0.204 / 0.113 | | | | | | | |

Change in functional connectivity was used as dependent variable, p-tau$_{181}$/Aβ$_{1-42}$ ratio at baseline, age at baseline, *APOE4* group, sex, education, time between baseline and FU24 assessment, and change in hippocampus volume from baseline to FU24 were used as independent variables. AD = Alzheimer's disease. FC = functional connectivity. PHC = parahippocampal cortex. r = right. PCun = precuneus. CI = 95% confidence interval. FU24 = follow-up 24 months after baseline. HC = hippocampus.

#### Table S27. Subsample A: Linear model of effects of change in hippocampal volume on memory change

| | Change in RBANS episodic memory performance over two years | | | | | | | | |
|---|---|---|---|---|---|---|---|---|---|
| *Predictors* | *Estimates* | *std. Error* | *std. Beta* | *standardized std. Error* | *CI* | *standardized CI* | *Statistic* | *p* | *std. p* |
| (Intercept) | -0.10 | 19.15 | -0.17 | 0.24 | -38.20 – 37.99 | -0.64 – 0.30 | -0.01 | 0.996 | 0.475 |
| Age in years | -0.03 | 0.19 | -0.02 | 0.12 | -0.42 – 0.35 | -0.25 – 0.21 | -0.18 | 0.861 | 0.861 |
| *APOE4* Group [carrier] | 1.08 | 2.39 | 0.21 | 0.24 | -3.68 – 5.84 | -0.27 – 0.69 | 0.45 | 0.653 | 0.383 |
| sex [female] | 1.13 | 2.05 | 0.14 | 0.26 | -2.95 – 5.21 | -0.38 – 0.66 | 0.55 | 0.583 | 0.583 |
| Education in years | -0.32 | 0.24 | -0.14 | 0.11 | -0.79 – 0.16 | -0.36 – 0.07 | -1.32 | 0.192 | 0.192 |
| Time baseline to FU24 | 0.01 | 0.02 | 0.08 | 0.11 | -0.02 – 0.05 | -0.14 – 0.30 | 0.70 | 0.484 | 0.484 |
| HC volume change over two years | 0.84 | 0.70 | 0.16 | 0.13 | -0.55 – 2.23 | -0.10 – 0.41 | 1.20 | 0.233 | 0.233 |

| | | | | | | | | |
|---|---|---|---|---|---|---|---|---|
| APOE4 carrier [carrier] × HC volume change over two years | -0.52 | 1.28 | -0.10 | 0.24 | -3.07 – 2.02 | -0.57 – 0.38 | -0.41 | 0.684 | 0.684 |
| Observations | 91 | | | | | | | | |
| $R^2$ / $R^2$ adjusted | 0.047 / -0.033 | | | | | | | | |

in amyloid and tau negative older adults

#### Table S28. Subsample A: Linear model of effects of connectivity and change in hippocampal volume on change in memory

| | Change in RBANS episodic memory performance over two years | | | | | | | |
|---|---|---|---|---|---|---|---|---|
| *Predictors* | *Estimates* | *std. Error* | *std. Beta* | *standardized std. Error* | *CI* | *standardized CI* | *Statistic* | *p* |
| (Intercept) | 2.93 | 18.72 | -0.19 | 0.23 | -34.31 – 40.17 | -0.65 – 0.26 | 0.16 | 0.876 |
| FC RSC r - PCC r at baseline | 8.53 | 4.04 | 0.23 | 0.11 | 0.48 – 16.57 | 0.01 – 0.45 | 2.11 | **0.038** |
| Age in years | -0.10 | 0.19 | -0.06 | 0.12 | -0.47 – 0.28 | -0.29 – 0.17 | -0.51 | 0.611 |
| *APOE4* Group [carrier] | 2.33 | 1.88 | 0.30 | 0.24 | -1.41 – 6.07 | -0.18 – 0.77 | 1.24 | 0.219 |
| sex [female] | 1.10 | 1.98 | 0.14 | 0.25 | -2.84 – 5.04 | -0.36 – 0.64 | 0.55 | 0.580 |
| Education in years | -0.26 | 0.23 | -0.12 | 0.11 | -0.73 – 0.21 | -0.33 – 0.09 | -1.11 | 0.272 |
| Time baseline to FU24 | 0.01 | 0.02 | 0.05 | 0.11 | -0.03 – 0.05 | -0.16 – 0.27 | 0.49 | 0.626 |
| HC volume change over two years | 0.62 | 0.58 | 0.12 | 0.11 | -0.52 – 1.77 | -0.10 – 0.33 | 1.09 | 0.281 |
| Observations | 91 | | | | | | | |
| $R^2$ / $R^2$ adjusted | 0.094 / 0.017 | | | | | | | |

Change in RBANS episodic memory performance in amyloid and tau negative older adults was used as dependent variable, functional connectivity at baseline, age at baseline, *APOE4* group, sex, education, time between baseline and FU24 assessment, and change in hippocampus volume from baseline to FU24 were used as independent variables. Due to multicollinearity, we left the interaction term of connectivity**APOE* out of the model. The interaction was not significant when included in the model. RBANS = Repeatable Battery for Assessment of Neuropsychological Status.

FC = functional connectivity. RSC = retrosplenial cortex. r = right. PCC = posterior cingulate cortex. CI = 95% confidence interval. FU24 = follow-up 24 months after baseline. HC = hippocampus.

#### Table S29. Subsample B: Linear model of effects of connectivity and change in hippocampal volume on change in memory

| | Change in RBANS episodic memory performance over two years | | | | | | | | |
|---|---|---|---|---|---|---|---|---|---|
| *Predictors* | *Estimates* | *std. Error* | *std. Beta* | *standardized std. Error* | *CI* | *standardized CI* | *Statistic* | *p* | *std. p* |
| (Intercept) | -10.54 | 25.08 | 0.01 | 0.26 | -60.74 – 39.67 | -0.51 – 0.53 | -0.42 | 0.676 | 0.982 |
| FC anterior HC r - superior PCun r at baseline | 9.26 | 10.64 | 0.14 | 0.16 | -12.04 – 30.56 | -0.18 – 0.47 | 0.87 | 0.388 | 0.388 |
| p-tau181/Aβ1-42 ratio at baseline | -43.00 | 63.51 | -0.10 | 0.15 | -170.13 – 84.13 | -0.41 – 0.20 | -0.68 | 0.501 | 0.501 |
| Age in years | 0.26 | 0.26 | 0.14 | 0.14 | -0.25 – 0.78 | -0.13 – 0.41 | 1.02 | 0.311 | 0.311 |
| *APOE4* Group [carrier] | 2.09 | 3.22 | 0.16 | 0.30 | -4.36 – 8.54 | -0.44 – 0.76 | 0.65 | 0.520 | 0.587 |
| sex [female] | -0.39 | 2.96 | -0.04 | 0.28 | -6.31 – 5.52 | -0.59 – 0.52 | -0.13 | 0.894 | 0.894 |
| Education in years | 0.20 | 0.48 | 0.06 | 0.13 | -0.76 – 1.16 | -0.21 – 0.32 | 0.42 | 0.679 | 0.679 |
| Time baseline to FU24 | -0.00 | 0.03 | -0.02 | 0.14 | -0.06 – 0.05 | -0.29 – 0.26 | -0.11 | 0.913 | 0.913 |
| HC volume change over two years | -0.50 | 0.75 | -0.09 | 0.14 | -2.00 – 1.00 | -0.37 – 0.19 | -0.67 | 0.508 | 0.508 |
| FC anterior HC r - superior PCun r at baseline × *APOE4* Group [carrier] | -43.12 | 16.97 | -0.66 | 0.26 | -77.09 – -9.16 | -1.18 – -0.14 | -2.54 | 0.014 | **0.014** |
| Observations | 68 | | | | | | | | |
| $R^2$ / $R^2$ adjusted | 0.136 / 0.002 | | | | | | | | |

Change in RBANS episodic memory performance was used as dependent variable, functional connectivity at baseline, p-tau181/Aβ1-42 ratio at baseline, age at baseline, *APOE4* group, sex, education, time between baseline and FU24 assessment, change in hippocampus volume from baseline to FU24, and the interaction term of connectivity**APOE* were used as independent variables. RBANS = Repeatable Battery for Assessment of Neuropsychological Status. FC = functional connectivity. HC = hippocampus. r = right. PCun = precuneus. CI = 95% confidence interval. FU24 = follow-up 24 months after baseline.

#### Table S30. Subsample A: Linear model of effects of change in connectivity, its interaction with *APOE* and change in hippocampal volume on change in memory

| | Change in RBANS episodic memory performance over two years | | | | | | | | |
|---|---|---|---|---|---|---|---|---|---|
| *Predictors* | *Estimates* | *std. Error* | *std. Beta* | *standardized std. Error* | *CI* | *standardized CI* | *Statistic* | *p* | *std. p* |
| (Intercept) | -2.00 | 19.22 | -0.19 | 0.24 | -40.24 – 36.24 | -0.66 – 0.28 | -0.10 | 0.917 | 0.435 |
| Change in FC RSC r - PCC r over two years | -0.50 | 3.91 | -0.02 | 0.13 | -8.29 – 7.28 | -0.27 – 0.24 | -0.13 | 0.898 | 0.898 |
| Age in years | -0.01 | 0.20 | -0.01 | 0.12 | -0.40 – 0.38 | -0.24 – 0.23 | -0.07 | 0.948 | 0.948 |
| *APOE4* Group [carrier] | 1.14 | 2.03 | 0.23 | 0.24 | -2.90 – 5.17 | -0.26 – 0.71 | 0.56 | 0.577 | 0.351 |
| sex [female] | 1.29 | 2.05 | 0.16 | 0.26 | -2.80 – 5.37 | -0.36 – 0.68 | 0.63 | 0.532 | 0.532 |
| Education in years | -0.29 | 0.24 | -0.13 | 0.11 | -0.78 – 0.19 | -0.36 – 0.09 | -1.21 | 0.231 | 0.231 |
| Time baseline to FU24 | 0.01 | 0.02 | 0.08 | 0.11 | -0.02 – 0.05 | -0.14 – 0.30 | 0.70 | 0.486 | 0.486 |
| HC volume change over two years | 0.76 | 0.61 | 0.14 | 0.11 | -0.45 – 1.96 | -0.08 – 0.37 | 1.25 | 0.216 | 0.216 |
| fc CG R 7 4-CG R 7 1 FU24>BL x *APOE4* Group [carrier] | -6.13 | 7.82 | -0.20 | 0.26 | -21.68 – 9.41 | -0.72 – 0.31 | -0.78 | 0.435 | 0.435 |
| Observations | 91 | | | | | | | | |
| $R^2$ / $R^2$ adjusted | 0.056 / -0.036 | | | | | | | | |

Change in RBANS episodic memory performance in amyloid and tau negative older adults was used as dependent variable, change in functional connectivity, age at baseline, *APOE4* group, sex, education, time between baseline and FU24 assessment, change in hippocampus volume from baseline to FU24, and the interaction of change in connectivity with *APOE4* group were used as independent variables. Due to multicollinearity, we left the interaction term of connectivity*age out of the model. The interaction was not significant when included in the model. RBANS =

Repeatable Battery for Assessment of Neuropsychological Status. FC = functional connectivity. RSC = retrosplenial cortex. r = right. PCC = posterior cingulate cortex. CI = 95% confidence interval. FU24 = follow-up 24 months after baseline. HC = hippocampus.

#### Table S31. Subsample B: Linear model of effects of change in connectivity and its interaction with *APOE* on change in memory

| | Change in RBANS episodic memory performance over two years | | | | | | | | |
|---|---|---|---|---|---|---|---|---|---|
| *Predictors* | *Estimates* | *std. Error* | *std. Beta* | *standardized std. Error* | *CI* | *standardized CI* | | *p* | *std. p* |
| (Intercept) | -10.60 | 26.60 | -0.05 | 0.27 | -63.85 – 42.66 | -0.60 – 0.50 | -0.40 | 0.692 | 0.863 |
| Change in FC anterior HC r - superior PCun over two years | -4.17 | 9.76 | -0.08 | 0.19 | -23.70 – 15.36 | -0.47 – 0.30 | -0.43 | 0.671 | 0.671 |
| p-tau$_{181}$/A$\beta_{1\text{-}42}$ ratio at baseline | -36.38 | 72.78 | -0.09 | 0.17 | -182.06 – 109.30 | -0.43 – 0.26 | -0.50 | 0.619 | 0.619 |
| Age in years | 0.25 | 0.27 | 0.13 | 0.14 | -0.29 – 0.80 | -0.16 – 0.42 | 0.92 | 0.361 | 0.361 |
| *APOE4* Group [carrier] | 1.09 | 3.34 | 0.05 | 0.31 | -5.61 – 7.78 | -0.56 – 0.66 | 0.33 | 0.746 | 0.874 |
| sex [female] | 0.14 | 3.11 | 0.01 | 0.29 | -6.10 – 6.37 | -0.57 – 0.59 | 0.04 | 0.966 | 0.966 |
| Education in years | 0.07 | 0.50 | 0.02 | 0.14 | -0.93 – 1.06 | -0.26 – 0.30 | 0.13 | 0.895 | 0.895 |
| Time baseline to FU24 | -0.00 | 0.03 | -0.00 | 0.15 | -0.06 – 0.06 | -0.30 – 0.29 | -0.01 | 0.992 | 0.992 |
| HC volume change over two years | -0.34 | 0.79 | -0.06 | 0.15 | -1.93 – 1.25 | -0.36 – 0.23 | -0.43 | 0.671 | 0.671 |
| Change in FC anterior HC r - superior PCun over two years × *APOE4* Group [carrier] | 16.19 | 14.18 | 0.32 | 0.28 | -12.19 – 44.56 | -0.24 – 0.88 | 1.14 | 0.258 | 0.258 |
| Observations | 68 | | | | | | | | |
| $R^2$ / $R^2$ adjusted | 0.052 / -0.095 | | | | | | | | |

Change in RBANS episodic memory performance was used as dependent variable, change in functional connectivity, p-tau$_{181}$/A$\beta_{1\text{-}42}$ ratio at baseline, age at baseline, *APOE4* group, sex, education, time between baseline and FU24 assessment, change in hippocampus volume from baseline to FU24, and the interaction of change in connectivity with

*APOE4* group were used as independent variables. Due to multicollinearity, we left the interaction term of connectivity* p-$tau_{181}$/$A\beta_{1-42}$ ratio at baseline out of the model. The interaction was not significant when included in the model. RBANS = Repeatable Battery for Assessment of Neuropsychological Status. FC = functional connectivity. HC = hippocampus. r = right. PCun = precuneus. CI = 95% confidence interval. FU24 = follow-up 24 months after baseline.
